# Transient receptor potential canonical 5 (TRPC5) mediates inflammatory mechanical pain

**DOI:** 10.1101/2020.07.13.201186

**Authors:** Katelyn E. Sadler, Francie Moehring, Stephanie I. Shiers, Lauren J. Laskowski, Alexander R. Mikesell, Zakary R. Plautz, Christina M. Mecca, Gregory Dussor, Theodore J. Price, John D. McCorvy, Cheryl L. Stucky

**Affiliations:** Department of Cell Biology, Neurobiology and Anatomy, Medical College of Wisconsin, Milwaukee, WI; University of Texas at Dallas, School of Behavioral and Brain Sciences and Center for Advanced Pain Studies, United States; Department of Anesthesiology, Medical College of Wisconsin, Milwaukee, WI

## Abstract

Persistent tactile pain is a poorly managed symptom of inflammatory and neuropathic injury. To develop therapies for this maladaptive sensation, the underlying molecular mediators must be identified. Using knockout mice and pharmacological inhibitors, we identified transient receptor canonical 5 (TRPC5) as a key contributor to the persistent tactile pain that occurs in many inflammatory and neuropathic preclinical rodent models. TRPC5 inhibition was effective in injuries associated with elevated levels of the bioactive phospholipid lysophosphatidylcholine (LPC). Exogenous application of LPC induced TRPC5-dependent behavioral mechanical allodynia, neuronal mechanical hypersensitivity, and spontaneous pain. *In vitro*, LPC activated both homomeric mouse and human TRPC5 channels, which upon examination of human dorsal root ganglia tissue, were expressed in 75% of human sensory neurons. Based on these results, TRPC5 inhibitors should be pursued as personalized therapy for spontaneous and tactile pain in conditions where elevated LPC is a biomarker.

## Main

The detection of non-painful (innocuous) touch and painful (noxious) pressure are critical for all organisms’ development, protection, and survival. However, after nerve damage or inflammatory injury, touch sensation can be distorted so that it is no longer beneficial. Instead, innocuous light touch is perceived as painful (i.e. mechanical allodynia) and thresholds for painful touch decrease *(1, 2)*. The molecular basis of these sensations is not understood, and therefore, managing touch-induced pain is difficult. A logical class of therapeutic targets is mechanically-sensitive ion channels. Indeed, Piezo2, a bonafide mechanically-gated innocuous touch transducer, and members of the Transient Receptor Potential (TRP) superfamily (e.g. TRPA1, TRPV4) contribute to mechanical hypersensitivity *(3–9)*. However, these channels are not ideal targets for further drug development considering they are also necessary for normal tactile sensation, proprioception, and pain sensation *(10–14)*. Instead, focus should be placed on proteins that are dispensable for normal tactile and pain sensation but recruited following injury to mediate persistent pain. Here, we identify transient receptor potential canonical 5 (TRPC5) as a novel tactile pain target. *In vivo*, TRPC5 is dispensable for normal innocuous and noxious tactile sensation in rodents *(15)*, and *in vitro*, TPRC5 channel activity is potentiated by inflammatory mediators *(16, 17)*. Despite these properties, TRPC5 has not been extensively explored as a therapeutic target for inflammatory pain. Using pharmacologic and genetic tools, we found that TRPC5 mediates tactile pain following inflammatory and neuropathic injury in mice. We identified lysophosphatidylcholine (LPC) as an endogenous agonist of both human and mouse TRPC5 channels, and found that TRPC5 inhibitor efficacy corresponded with tissue LPC levels following injury. To assess the translational potential of TRPC5 inhibitors, we also examined TRPC5 expression in human dorsal root ganglia neurons and found that TRPC5 was expressed in 75% of sensory neurons. Based on these data, LPC can be used as a biomarker for the tailored application of TRPC5 inhibitors, which have potential as non-opioid based therapies for tactile and spontaneous pain.

## Results

### TRPC5 activity is required for behavioral mechanical hypersensitivity in persistent pain models

To assess the role of TRPC5 in persistent mechanical allodynia, a battery of inducible pain models was performed in wildtype and global TRPC5 knockout (C5KO) mice (**Fig 1A**). First, intraplantar injection of Complete Freund’s Adjuvant (CFA) was used to induce severe inflammatory pain. C5KO mice and wildtype controls developed similar levels of mechanical allodynia immediately (2 hr) following CFA injection, but C5KO mice began to lose this hypersensitivity 24 hours following injection. Two days after CFA injection, C5KO mice no longer exhibited the mechanical allodynia that persisted for > 40 days in wildtype mice (**Fig 1B**). C5KO mice also failed to exhibit hypersensitivity to noxious punctate stimuli (**Fig 1C**) or dynamic mechanical allodynia (**Fig 1D**) following CFA injection, unlike wildtype controls. To model clinical application, we assessed the analgesic efficacy of AC1903, a selective small molecule TRPC5 inhibitor *(18)*, in established CFA-induced mechanical allodynia. AC1903 eliminated mechanical allodynia in wildtype mice when injected two, seven, or 21 days following CFA injection (**Fig 1E**). Importantly, AC1903 did not change the mechanical sensitivity of uninjured wildtype mice (**Fig 1E, 1F)** or injured C5KO mice (**Fig 1F**), suggesting that the effects of AC1903 are specific to TRPC5 channel activity, which is not involved in naïve mechanosensation *(15)*.

**Fig 1.**
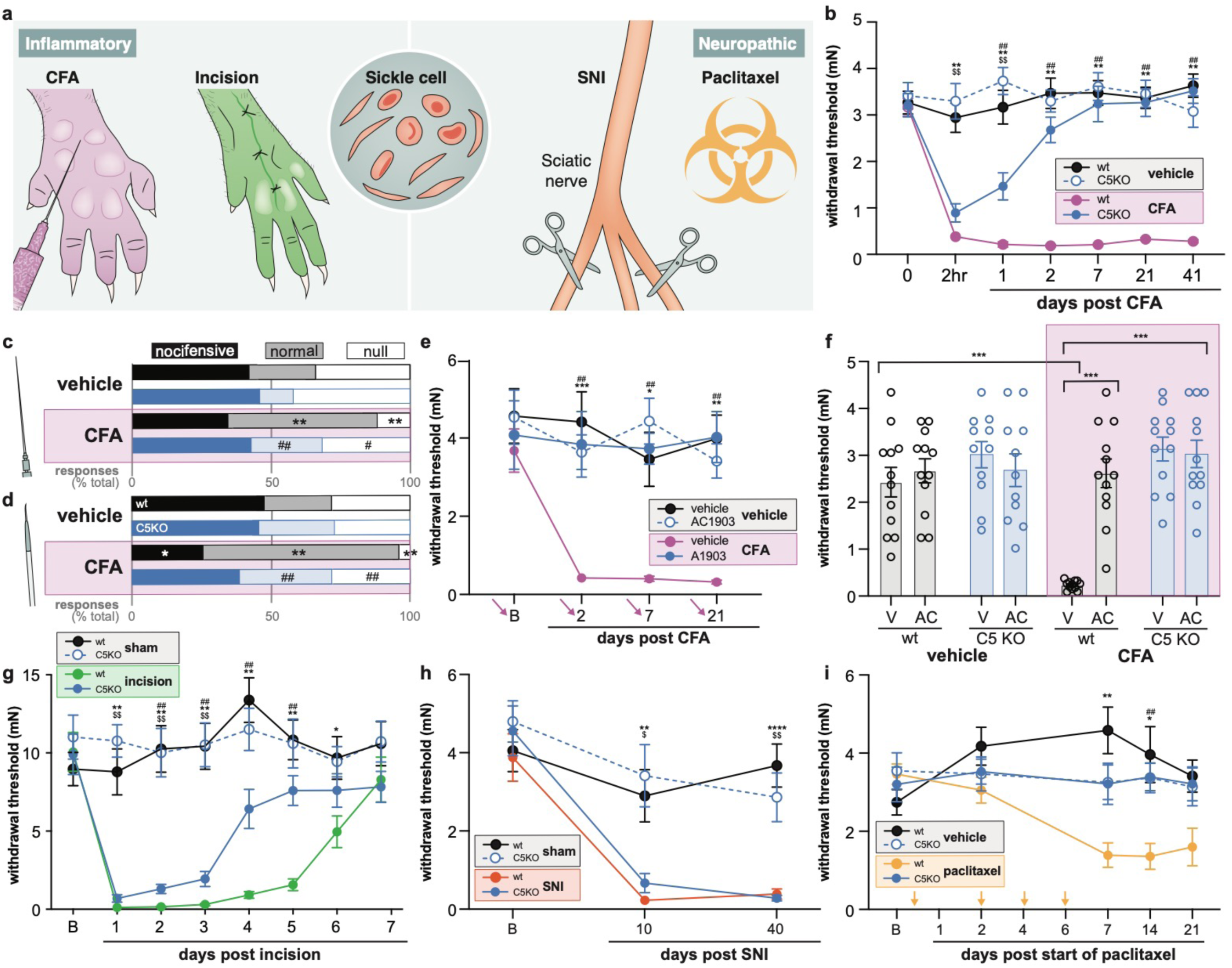
TRPC5 contributes to persistent mechanical hypersensitivity in inducible inflammatory pain models. **a**. Depiction of common rodent pain models. Complete Freund’s Adjuvant (CFA) injection and hindpaw plantar incision are inflammatory pain models; spared nerve injury (SNI) and paclitaxel-induced peripheral neuropathy are neuropathic. Transgenic sickle cell disease (SCD) mice exhibit inflammatory and neuropathic pain properties. **b**. Mechanical withdrawal thresholds of global TRPC5 knockout (C5KO) and wildtype (wt) mice following CFA injection; n=11-12 mice. **c**. Response characterization to noxious needle hindpaw stimulation 9 days post-CFA; n=6-10 mice; bars are group averages. **d**. Response characterization to paintbrush swiping across plantar hindpaw 9 days post-CFA; n=6-10 mice; bars are group averages. **e**. Mechanical withdrawal thresholds 60 min following intraplantar injection of AC1903; n=10; arrows indicate injection on each day. **f**. Mechanical withdrawal thresholds on day 41 post-CFA, 60 min following intraplantar injection of AC1903 (AC) or vehicle (V); n=11-12; Bonferroni post-hoc comparison \*\*\**P*<0.001. **g**. Mechanical withdrawal thresholds following hindpaw plantar incision; n=17-21. **h**. Mechanical withdrawal thresholds following SNI; n=7-10. **i**. Mechanical withdrawal thresholds following paclitaxel; n=8-10; arrows indicate paclitaxel injections. Post-hoc comparisons (Bonferroni or Fisher’s exact) for all panels unless otherwise stated.: \**P*<0.05, \*\**P*<0.01, \*\*\**P*<0.001 wildtype control vs. injury; ^$^*P*<0.05, ^$$^*P*<0.01, ^$$$^*P*<0.001 C5KO control vs. injury; ^#^*P*<0.05, ^##^*P*<0.01, ^###^*P*<0.001 injury C5KO vs. wildtype. All data are mean ± SEM unless otherwise stated; B: baseline; AC1903 dose: 50 µg.

While the CFA model is both reliable and robust, CFA injections lack face validity and have low translational potential. Therefore, we utilized the hindpaw incision model of post-surgical pain *(19)* to determine if TRPC5 activity contributes to clinically-relevant inflammatory pain. Like in the CFA model, C5KO and wildtype mice exhibited similar levels of mechanical allodynia shortly after incision (**Fig 1G**). However, C5KO mechanical thresholds returned to sham levels by postoperative day 5, whereas wildtype thresholds did not recover until day 7. On postoperative day 4, incised C5KO mice did not exhibit the hypersensitivity to noxious punctate stimuli (**Fig S1A**) or dynamic brush allodynia (**Fig S1B**) that was observed in incised wildtype mice. Similar to established CFA mechanical allodynia, established postoperative mechanical allodynia in wildtype mice was partially alleviated by AC1903 treatment (**Fig S1C**). Based on these collective data, TRPC5 activity contributes to the persistent mechanical hypersensitivity that occurs following peripheral inflammatory tissue injuries. The reduced mechanical allodynia observed in C5KO mice did not result from decreased peripheral inflammation because wildtype and C5KO mice exhibited similar levels of hindpaw swelling following CFA injections (**Fig S1D)** and hindpaw incision (**Fig S1E**). Therefore, TRPC5 inhibitors could be pursued as therapeutics that specifically decrease the maladaptive sensations associated with many persistent inflammatory conditions.

To determine whether TRPC5 is also required for neuropathic mechanical allodynia, spared nerve injury (SNI) and chemotherapy induced peripheral neuropathy (CIPN) models were employed in wildtype and C5KO mice. Unlike in the inflammatory models, C5KO and wildtype mice that underwent SNI exhibited similar levels of mechanical allodynia (**Fig 1H**), hypersensitivity to noxious punctate stimuli (**Fig S1F**), and dynamic brush allodynia (**Fig S1G**). Peripheral AC1903 administration was also ineffective in alleviating SNI-induced mechanical allodynia (**Fig S1H**). The lack of TRPC5 involvement in this traumatic nerve injury model starkly contrasted to channel contributions in inflammatory pain models. To determine if this was unique to SNI or more broadly applicable to other neuropathic conditions, C5KO and wildtype mice were treated with paclitaxel, a neurotoxic chemotherapeutic agent. C5KO mice failed to develop mechanical allodynia (**Fig 1I**), hypersensitivity to noxious punctate stimuli (**Fig S1I**), or dynamic brush allodynia (**Fig S1J**) following paclitaxel treatment, unlike wildtype controls which exhibited all three symptoms.

### The endogenous lipid lysophosphatidylcholine (LPC) induces TRPC5-dependent mechanical hypersensitivity following injury

One potential site of action for TRPC5-related analgesia are peripheral sensory neurons of the dorsal root *(15)* and trigeminal ganglia *(20)*. Since TRPC5 is indirectly activated by stretch *(21)*, injury-specific sensitization of this channel could increase the mechanical sensitivity of peripheral sensory neurons, and thereby increase behavioral responses to innocuous and noxious touch. Mechanical hypersensitivity is well documented in C fibers following CFA *(8, 22)*, hindpaw incision *(23)*, SNI *(24)*, and paclitaxel *(25)* injury, but it is unclear if this hypersensitivity is mediated by TRPC5. To specifically assess if peripheral TRPC5 contributes to neuronal mechanical hypersensitivity, *ex vivo* recordings of tibial nerve C fibers were performed in wildtype and C5KO tissues isolated 7 days following CFA injection (**Fig 2A**). Compared to C fibers from wildtype CFA-injected mice, afferents from C5KO CFA-injected mice had elevated mechanical thresholds (**Fig 2B, 2C**) and reduced firing during dermal force application (**Fig 2D, 2E**). Mechanical thresholds (**Fig S2A)** and mechanically-induced firing frequencies (**Fig S2B)** were similar in C fibers isolated from vehicle-treated wildtype and C5KO mice. These *ex vivo* experiments suggest that peripherally-expressed TRPC5 contributes to the persistent mechanical hypersensitivity observed in the CFA model, likely through sensitization or activation of the channel via a chemical mediator present in the injured tissue. In order to identify potential mediators, we performed mass spectrometry on the tissues damaged in each of the injury models. Lysophosphatidylcholine (LPC; **Fig 2F**) *(26)*, a pro-inflammatory single-chain fatty acid *(27)*, was markedly elevated in the skin of both wildtype and C5KO mice following intraplantar CFA injections (**Fig 2G**). Increases were observed in LPC species with varying carbon tail lengths and double bond number (**Table 1**). Circulating LPC levels were unchanged in CFA-injected animals (**Fig S3A; Table 2**), suggesting that the excess lipid is derived from cells localized to the injured tissue. Similar injury site-specific increases in LPC were also found in the incised hindpaw skin of wildtype and C5KO mice (**Fig 2H, Tables 1 and 2, Fig S3B**). In contrast to the inflammatory models, total LPC levels and individual species quantities were unchanged in the sciatic nerve (**Fig 2I, Tables 3**) or serum (**Fig S3C, Table 2**) of wildtype or C5KO mice following SNI, perhaps explaining the lack of efficacy of TRPC5 inhibition or deletion in this injury model.

**Table 1.** Relative amount of LPC species in skin 5 days post-hindpaw plantar incision or 7 days post-CFA. Values are fold increases relative to wildtype (wt) sham/vehicle. Red italic text: *P*<0.05 vs. wt sham; blue bold text: *P*<0.01 vs. wt sham; green underlined text *P*<0.001 vs. wt sham; pink cell fill: *P*<0.05 vs. C5KO sham; blue cell fill: *P*<0.01 vs. C5KO sham; green cell fill: *P*<0.001 vs. C5KO sham; nd: not detected in any sample; x: not detected in wt sham; n=3 mice.

|  | INCISION |  |  | CFA |  |  |
| --- | --- | --- | --- | --- | --- | --- |
|  | incision | C5KO sham | C5KO incision | CFA | C5KO vehicle | C5KO CFA |
| (15:0) | *1.81* | 0.44 | 1.63 | *1.68* | 0.76 | **1.53** |
| (16:0) | *1.52* | 1.07 | 1.49 | <u>2.03</u> | 0.99 | <u>2.21</u> |
| (16:1) | 1.35 | 0.96 | 1.19 | 1.73 | 0.95 | 1.73 |
| (17:0) | 1.02 | 0.89 | 1.12 | *1.52* | 0.88 | *1.53* |
| (18:0) | *1.15* | 0.81 | 1.09 | <u>1.38</u> | 0.89 | <u>1.45</u> |
| (18:1) | 1.39 | 0.93 | 1.30 | **2.04** | 0.98 | *2.06* |
| (18:2) | 1.25 | 1.04 | **1.24** | 1.81 | 0.90 | 1.78 |
| (18:3) | 1.39 | 0.98 | 1.65 | 1.81 | 1.02 | 1.73 |
| (20:0) | 2.17 | nd | 5.53 | nd | nd | x |
| (20:1) | 1.16 | 0.87 | 0.95 | 1.31 | 0.80 | 1.14 |
| (20:2) | x | x | x | *x* | nd | **x** |
| (20:3) | 1.36 | 0.84 | 1.71 | **1.94** | 1.06 | *2.32* |
| (20:4) | 1.26 | 1.09 | 1.17 | *2.14* | 1.13 | *1.82* |
| (20:5) | 1.40 | 0.77 | 1.23 | 1.88 | 0.97 | 1.85 |
| (22:4) | nd | nd | nd | nd | nd | nd |
| (22:5) | 0.87 | 1.16 | nd | 0.97 | 1.03 | nd |
| (22:6) | nd | nd | nd | x | nd | nd |

**Table 2.** Relative amount of serum LPC species in Berk SS sickle cell disease mice, 5 days post-hindpaw plantar incision, 10 days post-SNI, or 7 days post-CFA. Values are fold increases relative to wildtype (wt) sham/vehicle. Red italic text: *P*<0.05 vs. wt sham; blue bold text: *P*<0.01 vs. wt sham; pink cell fill: *P*<0.05 vs. C5KO sham; nd: not detected in any sample; x: not detected in wt sham; n=3 mice.

|  | Berk SS | INCISION |  |  | SNI |  |  | CFA |  |  |
| --- | --- | --- | --- | --- | --- | --- | --- | --- | --- | --- |
|  |  | incision | C5KO sham | C5KO incision | SNI | C5KO sham | C5KO SNI | CFA | C5KO vehicle | C5KO CFA |
| (15:0) | 0.95 | 0.93 | 0.76 | 0.94 | 0.83 | 0.89 | 0.99 | 0.38 | 1.74 | 0.86 |
| (16:0) | *1.15* | 0.94 | 0.97 | 1.05 | 1.10 | 0.83 | 1.01 | 0.86 | 1.20 | 0.95 |
| (16:1) | **2.17** | **0.68** | 0.92 | 0.86 | 1.12 | 0.88 | 0.94 | 0.87 | 1.08 | 0.93 |
| (17:0) | 0.87 | 0.89 | 0.85 | 1.09 | 1.13 | 1.07 | 1.24 | 0.77 | 1.31 | 0.93 |
| (18:0) | 0.96 | 0.91 | 0.96 | 1.17 | 1.14 | 0.78 | 0.97 | 0.95 | 1.23 | 1.10 |
| (18:1) | **2.12** | 0.79 | 0.96 | 0.77 | 1.09 | 0.90 | 0.85 | 0.89 | 1.04 | 0.94 |
| (18:2) | *1.39* | 0.84 | 0.83 | 1.01 | 1.00 | 0.95 | 0.85 | 0.95 | 1.17 | 0.97 |
| (18:3) | 2.08 | nd | 1.13 | 0.92 | nd | 1.18 | nd | 0.58 | 0.81 | 0.70 |
| (20:0) | **0.39** | 1.09 | 1.17 | 1.08 | 1.32 | 1.44 | *3.12* | 1.23 | 1.29 | 0.98 |
| (20:1) | 0.90 | 1.04 | 1.24 | 0.99 | 1.49 | 1.14 | 1.63 | 0.80 | 1.09 | 0.88 |
| (20:2) | **1.79** | 0.88 | 0.94 | *0.72* | 1.13 | 0.97 | 1.25 | 0.95 | 0.95 | 1.06 |
| (20:3) | **1.77** | 0.84 | 1.00 | 0.77 | 1.08 | 0.96 | 0.97 | 1.08 | 1.03 | 1.06 |
| (20:4) | **2.83** | 0.82 | 0.69 | 1.05 | 0.94 | 0.71 | 0.62 | 0.82 | 1.20 | 0.78 |
| (20:5) | nd | nd | nd | nd | nd | nd | nd | nd | nd | x |
| (22:4) | **7.34** | 1.13 | 0.96 | 0.84 | 1.10 | 0.82 | 1.05 | 0.75 | 0.59 | 0.89 |
| (22:5) | *1.81* | *0.68* | **0.59** | **0.54** | 1.19 | 0.62 | 1.04 | 0.81 | 0.85 | 0.84 |
| (22:6) | **2.05** | 0.85 | 1.06 | *1.34* | 1.17 | 1.15 | 0.94 | 0.83 | 1.33 | 0.96 |

**Table 3.** Relative amount of LPC species in sciatic nerve 10 days post-SNI; nd: not detected in any sample; x: not detected in wt sham; n=3 mice.

|  | SNI |  |  |
| --- | --- | --- | --- |
|  | SNI | C5KO sham | C5KO SNI |
| (15:0) | nd | nd | nd |
| (16:0) | 0.75 | 0.50 | 0.68 |
| (16:1) | nd | 0.64 | 0.77 |
| (17:0) | nd | nd | x |
| (18:0) | 0.67 | 0.44 | 0.70 |
| (18:1) | 0.70 | 0.53 | 0.50 |
| (18:2) | 0.59 | 0.44 | 0.85 |
| (18:3) | nd | nd | nd |
| (20:0) | nd | nd | nd |
| (20:1) | nd | nd | nd |
| (20:2) | nd | nd | nd |
| (20:3) | nd | nd | nd |
| (20:4) | nd | nd | nd |
| (20:5) | nd | nd | nd |
| (22:4) | nd | nd | nd |
| (22:5) | nd | nd | nd |
| (22:6) | nd | nd | nd |

**Fig 2.**
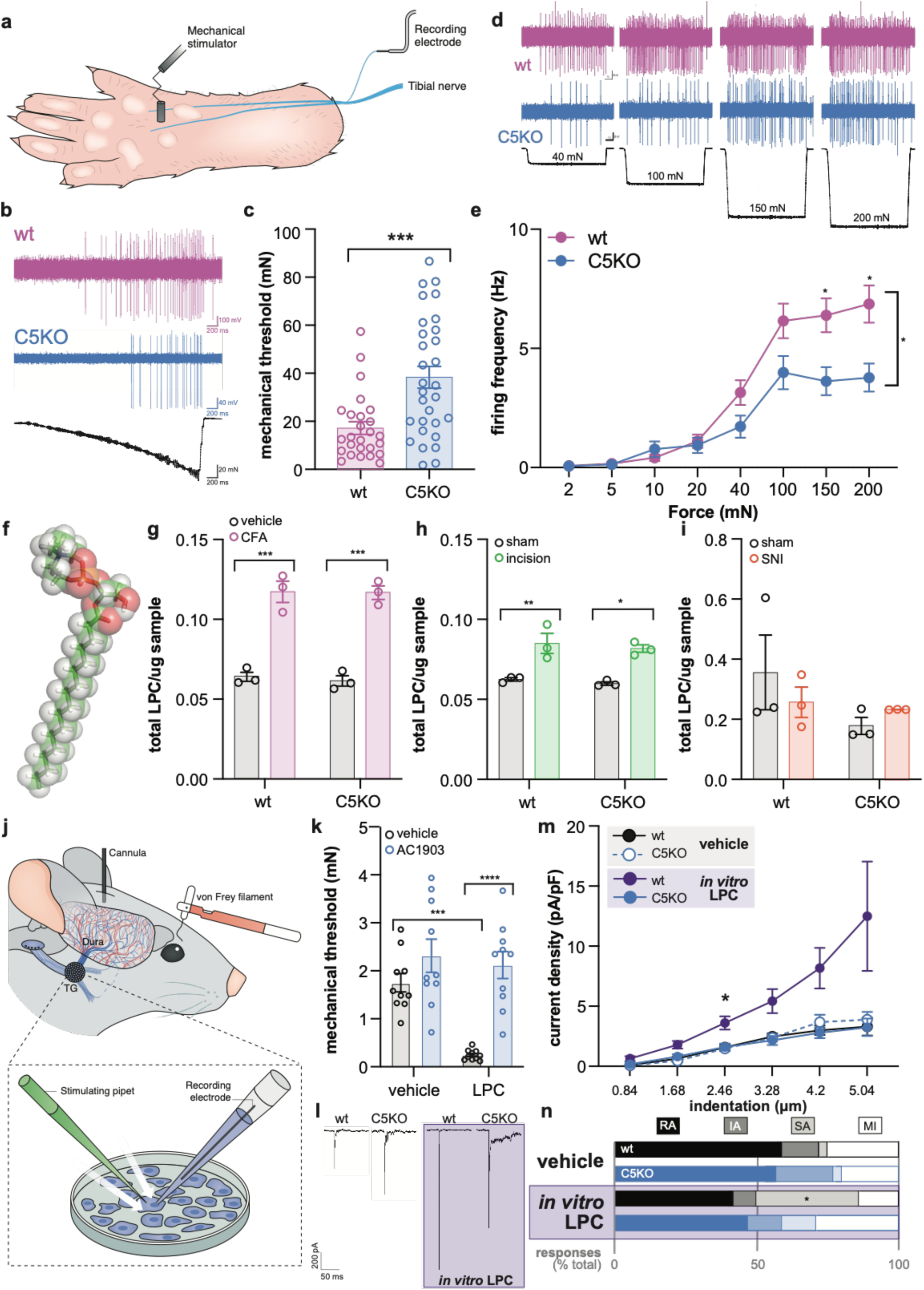
Injury-related increases in lysophosphatidylcholine (LPC) induce mechanical hypersensitivity. **a**. Schematic of teased fiber recordings. **b**. Representative C fiber activity traces during force ramp application (10 s; up to 100 mN). Force ramps were used to determine the mechanical threshold of each fiber. **c**. Mechanical thresholds of fibers from CFA-injected wt and C5KO mice; n=27-30 units from 5 mice; Mann Whitney test \*\*\**P*<0.001. **d**. Representative C fiber activity during static force application. **e**. Mean mechanically-induced firing rates of C fibers from CFA-injected mice; n=27-30 units from 5 mice, two-way ANOVA effect of genotype \**P*<0.05. **f**. 3D model of LPC (16:0). **g**. Relative amount of LPC in skin 7 days post-CFA injection; n=3 mice. **h**. Relative amount of LPC in skin 5 days post-hindpaw plantar incision; n=3 mice. **i**. Relative amount of LPC in sciatic nerve 10 days post-SNI; n=3 mice. **j**. Schematic of dural injection, facial von Frey testing, and focal mechanical stimulation of isolated sensory neurons. **k**. Periorbital mechanical withdrawal thresholds 60 min following dural application of 4.8 nM LPC and/or 50 µg of AC1903; n=10. **l**. Representative current traces from trigeminal ganglia (TG) neurons during mechanical stimulation. **m**. Mean mechanical current density of TG neurons during 250 nM LPC incubation; n=24-30 neurons from 3-5 mice. **n**. Characterization of TG mechanical currents as rapidly (RA), intermediate (IA), slowly (SA) adapting, or mechanically insensitive (MI); bars are group averages. Bonferroni post-hoc comparisons for all panels: \**P*<0.05, \*\**P*<0.01, \*\*\**P*<0.001, \*\*\*\**P*<0.0001. All data are mean ± SEM unless otherwise stated.

After observing injury-induced increases in LPC, we next wanted to determine if LPC directly contributed to mechanical allodynia. To do this, we administered LPC onto the dura mater of wildtype mice. Dural injections have been used to induce migraine-like pain in rodents *(28)* (**Fig 2J**). Notably, patients with migraine exhibit mechanical allodynia *(29)* and have higher levels of circulating LPC species *(30)*, but a direct relationship between lipid levels and mechanical allodynia has not yet been reported. In wildtype mice, dural infusion of LPC increased the mechanical sensitivity of the periorbital region (**Fig S4A**). However, when LPC was co-infused with AC1903, animals did not develop mechanical allodynia (**Fig 2K**). To determine if this behavioral hypersensitivity resulted from direct neuronal sensitization, we performed whole-cell patch clamp recordings on trigeminal ganglia (TG) neurons exposed to LPC (**Fig 2J, 2L**). LPC increased the amplitude of mechanically-evoked whole-cell currents in wildtype neurons, but had no effect on the mechanically-evoked current amplitude of C5KO neurons (**Fig 2M**). Further, LPC incubation altered the mechanically-evoked current profiles of wildtype neurons; more slowly adapting (SA) currents were observed when the recording bath contained LPC (**Fig 2N**). LPC incubation did not change the mechanical current types observed in C5KO neurons. To summarize, local release of LPC during injury-associated tissue inflammation sensitizes primary afferent responses to mechanical stimulation in a TRPC5-dependent manner. Based on these findings, TRPC5 inhibitors would effectively alleviate persistent mechanical allodynia in injuries that are characterized by increases in LPC.

### LPC directly activates murine TRPC5 and induces behavioral aversion

As illustrated in **Figure 2**, direct application of LPC induced neuronal sensitivity to mechanical probing. However, it was unclear whether this increased sensitivity resulted from LPC directly activating TRPC5, or indirectly modulating channel function. To answer this question, LPC-induced calcium flux was measured in HEK cells stably transfected with mouse TRPC5 (mTRPC5). LPC specifically activated mTRPC5 when it was applied up to 32 µM (**Fig 3A, 3B, Fig S5A**). However, when LPC concentrations exceeded 100 µM, non-specific calcium flux was noted in both transfected and non-TRPC5 transfected HEK cells, likely due to detergent-like properties of LPC at high concentrations (**Fig 3A**). Based on these data, it is likely that inflammation-related pain partially arises as a result of LPC effects at TRPC5. Since AC1903 decreased these pain-like behaviors in mice, we next asked if AC1903 and other TRPC5 inhibitors that directly bind to TRPC5 *(31)* act by selectively blocking LPC activity at the channel. AC1903, HC-070 *(32)*, and ML204 *(33)*, decreased LPC-induced calcium flux through mTPRC5 (**Fig 3B**). These compounds were similarly effective in blocking mTRPC5 activation by (-)-Englerin A, a more TRPC-selective channel agonist (**Fig S5B**). *In vitro* potency of these inhibitors roughly translated to *in vivo* analgesia; HC-070 and AC1903 were similarly effective in reducing hindpaw incision mechanical allodynia, but ML204 had no effect on mechanical thresholds (**Fig 3C**).

**Fig 3.**
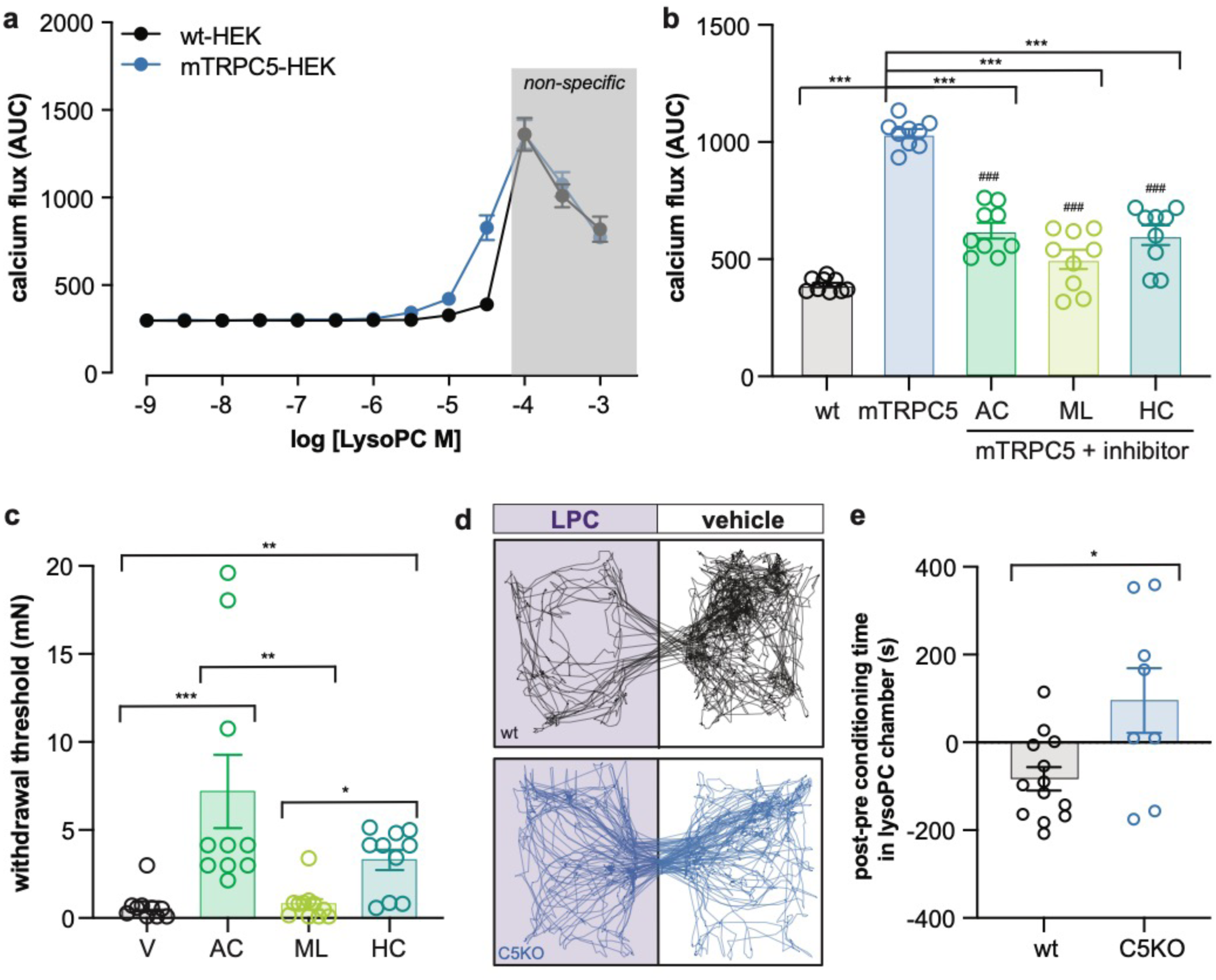
LPC directly activates mouse TRPC5 (mTRPC5). **a**. LPC-mediated calcium flux in HEK cells un-transfected or stably expressing mTRPC5; n=3 independent replicates performed in triplicate. **b**. Quantification of LPC (32 µM)-induced calcium flux in HEK cells un-transfected or stably expressing mTRPC5, and incubated with ML204 (320 µM), HC-070 (100 µM), or AC 1903 (320 µM); n=3 or more independent replicates performed in triplicate; Bonferroni post-hoc ^###^*P*<0.001 wt vs. inhibitor. **c**. Mechanical withdrawal thresholds on day 4 post-hindpaw plantar incision, 60 min following intraplantar injection of PBS vehicle (V), AC1903 (50 µg), ML204 (50 µg), or HC-070 (50 ng); n=10 mice. **d**. Representative track plots of wildtype (wt) and TRPC5 knockout (C5KO) mice in two-chamber box during post-conditioning test; LPC injection (5 µM) associated with purple box, vehicle injection associated with white box. **e**. Difference in LPC chamber time between pre- and post-conditioning trials; n=8-13 mice; unpaired t-test \**P*<0.05. Bonferroni post-hoc comparisons for all panels: \**P*<0.05, \*\**P*<0.01, \*\*\**P*<0.001, \*\*\*\**P*<0.0001. All data are mean ± SEM.

Since LPC is capable of directly activating TRPC5, we hypothesized that this lipid may also induce spontaneous pain following injury. To assess this behaviorally, we performed conditioned place aversion experiments in wildtype and C5KO mice. In a two-chamber conditioning paradigm, wildtype mice developed a place aversion for a chamber associated with 5 µM LPC treatment (**Fig 3D**), but not 1 µM (**Fig S4B**). C5KO mice did not develop the same place aversion as wildtype mice following one (**Fig S4C**) or three days of LPC conditioning (**Fig 3E**). Based on these data, TRPC5 modulators may also be effective therapies for spontaneous pain.

### TRPC5-mediated mechanical hypersensitivity persists after removal of extracellular LPC

Up to this point, our data demonstrate that direct neuronal exposure to LPC induces TRPC5-dependent increases in mechanical sensitivity. To determine if TRPC5-dependent hypersensitivity persists in the absence of extracellular LPC, we measured the mechanical sensitivity of TG neurons 48 hr following *in vivo* application of LPC. Neurons were isolated from wildtype or C5KO mice 24 hr after dural application of LPC, and 24 hr later, mechanical sensitivity was assessed using whole-cell patch clamp recordings. Neurons from LPC-injected wildtype mice exhibited larger mechanical currents than neurons isolated from LPC-injected C5KO mice or vehicle-injected wildtype animals (**Fig 4A**). Similar sensitization was also observed in a behavioral model of migraine-like pain. Migraine attacks can be triggered by subthreshold innocuous stimuli at times when patients are otherwise normal. To determine if LPC-induced TRPC5 activation and subsequent sensitization in the absence of the lipid contribute to this phenomenon, LPC dural injections were again performed in wildtype and C5KO mice. Similar to **Figure 2K**, LPC application induced mechanical allodynia in wildtype mice. After mechanical thresholds recovered however, subsequent infusion of pH 7.0 synthetic interstitial fluid on the dura induced facial mechanical allodynia in wildtype animals that had previously received an LPC infusion (**Fig 4B**). C5KO mice did not develop the initial LPC-induced mechanical allodynia or subsequent pH 7.0-induced hypersensitivity. These data suggest that LPC activation of TRPC5 may be sufficient to induce neuronal and behavioral sensitization that persists even in the absence of the endogenous ligand.

**Fig 4.**
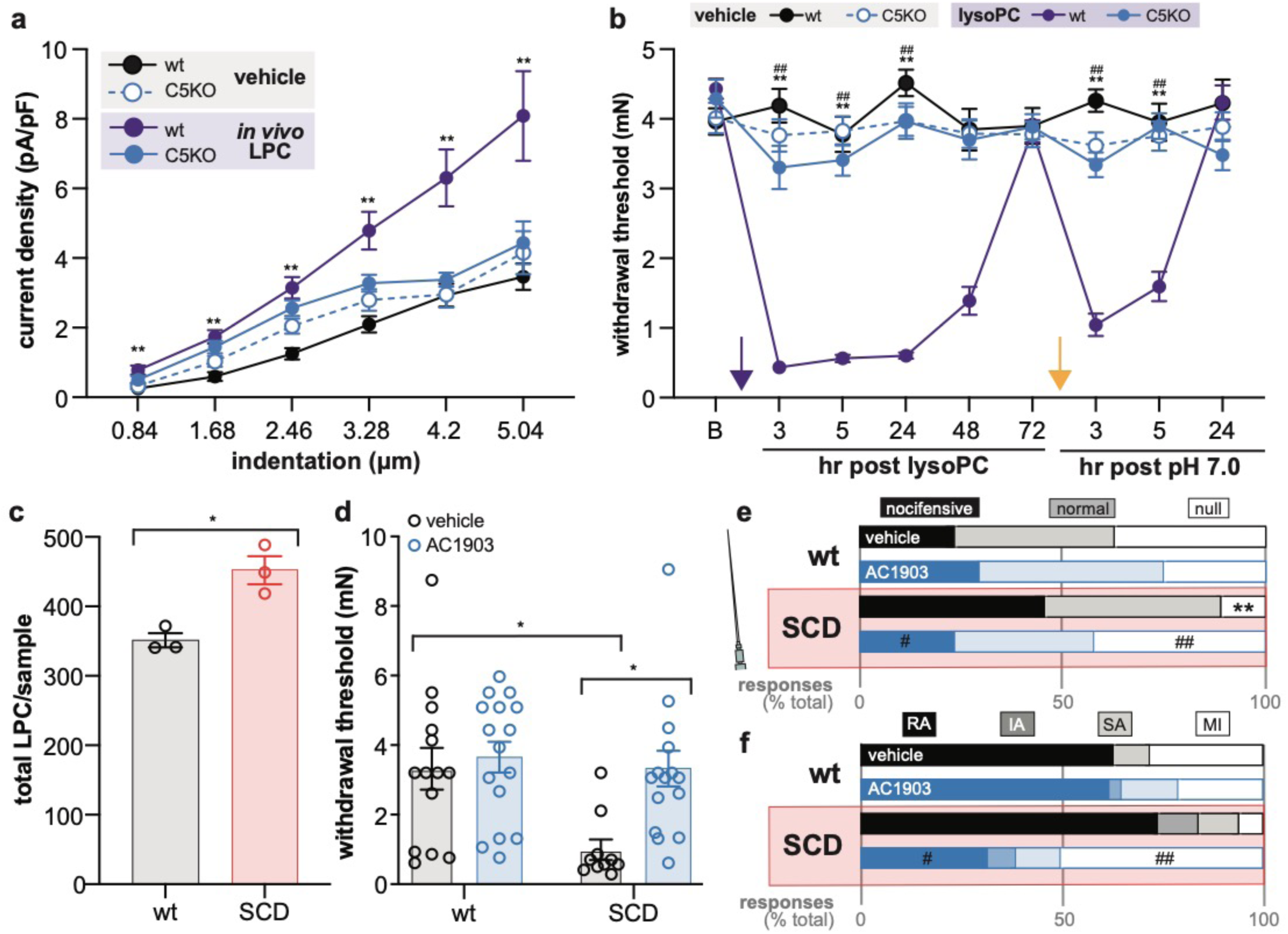
After initial exposure to LPC, TRPC5 activity mediates mechanical hypersensitivity even in the absence of LPC. **a**. Mean mechanical current density of TG neurons isolated from mice 24 hr following dural application of 4.8 nM LPC; n=33-45 cells from 3-5 mice, Bonferroni post-hoc comparison \*\**P*<0.01 wildtype vehicle vs. LPC. **b**. Periorbital mechanical withdrawal thresholds following dural application of 1.2 nM LPC and pH 7.0 PBS; n=10-15; purple arrow indicates LPC injection, yellow arrow indicates pH 7.0 PBS injection; Bonferroni post-hoc comparison \*\**P*<0.01 wildtype vehicle vs. LPC, ^##^ *P*<0.01 LPC wildtype vs. C5KO. **c**. Relative amount of LPC in serum of wildtype (wt) and sickle cell disease (SCD) mice; n=3 mice, unpaired t-test \**P*<0.05. **d**. Mechanical withdrawal thresholds 60 min following intraplantar injection of AC1903 (50 µg); n=10-14 mice, Bonferroni post-hoc comparisons \**P*<0.05. **e**. Response characterization to noxious needle hindpaw stimulation 60 min following intraplantar injection of AC1903 (50 µg); n=9-10 mice, bars are group averages, Fishers exact test \*\**P*<0.01 vehicle wildtype vs. SCD, ^#^*P*<0.05, ^##^*P*<0.01 SCD vehicle vs. AC1903. **f**. Characterization of DRG mechanical currents as rapidly (RA), intermediate (IA), slowly (SA) adapting, or mechanically insensitive (MI); bars are group averages, Fishers exact test ^#^*P*<0.05, ^##^*P*<0.01 SCD vehicle vs. AC1903. All data are mean ± SEM unless otherwise stated.

This idea was further confirmed in a transgenic model of sickle cell disease (SCD). SCD mice have higher levels of circulating LPC than controls (**Fig 4C**), likely from disease-related changes in erythrocyte membrane lipid composition *(34)*. AC1903 treatment significantly decreased the behavioral mechanical allodynia (**Fig 4D**) and hypersensitivity to noxious mechanical stimuli (**Fig 4E**) observed in SCD mice, paralleling AC1903 efficacy in other models characterized by increased LPC. When dorsal root ganglia neurons were isolated from the LPC-rich native environment however, AC1903 application still decreased the mechanical sensitivity of these cells. AC1903 incubation increased the number of SCD neurons that were mechanically insensitive in whole-cell patch clamp experiments, whereas the compound had no effect on neurons isolated from wildtype animals (**Fig 4F**). These data suggest that increased TRPC5 activity may persist even in the apparent absence of an endogenous agonist. TRPC5 inhibitors may therefore be effective, non-opioid based chronic pain therapies.

### TRPC5 is expressed in human sensory neurons and activated by LPC

The analgesic efficacy of TRPC5 inhibitors across seemingly disparate mouse pain models provides rationale for advancing this target to clinical trials. However, it would be remiss to not acknowledge the history of failed targets that were originally characterized as promising analgesics using these exact same animal models *(35)*. One potential reason for this limited translational success is species-specific differences in target expression. To this end, we performed RNAscope *in situ* hybridization to characterize TRPC5 expression in human DRG neurons (**Fig 5A, Table 4**). TRPC5 expression was observed in 75% of human DRG neurons (**Fig 5B**), particularly enriched in medium and small diameter neurons (cell bodies <80 µm; roughly equivalent to <40 µm diameter cell bodies in mouse (**Fig 5C**)). Of the neurons expressing TRPC5, ∼80% could be classified as nociceptors, co-expressing either CGRP (*CALCA*) or P2×3R (*P2RX3*) (**Fig 5D**). The remaining 20% of TRPC5-positive neurons did not express either nociceptor marker and are likely part of the Aβ-mechanoreceptor population *(36)*. This TRPC5 expression pattern in human DRG neurons is almost identical to that observed for TRPV1 *(36)*, another member of the TRP channel superfamily that has been a primary focus of both preclinical and clinical pain studies for the last two decades. In addition to verifying expression, we also assessed if human TRPC5 (hTRPC5) channels were activated by LPC. Like mTRPC5, hTRPC5 activity was specifically induced by physiologically relevant concentrations of LPC (**Fig 5E, 5F, Fig S5C**); non-specific, possibly detergent-mediated effects were observed when >100 µM concentration of LPC was applied to cells. Importantly, application of AC1903, HC-070, and ML204 effectively blocked both LPC (**Fig 5F**) and (-)-Englerin A (**Fig S5D**), induced activation of hTRPC5. These experiments support the use of TRPC5 inhibitors in persistent pain conditions, specifically those that are characterized by elevated levels of LPC.

**Table 4.** Human DRG tissue information.

| Donor # | DRG | Sex | Age | Race | Surgery/Cause of Death |
| --- | --- | --- | --- | --- | --- |
| 1 | L5 | Male | 22 | Hispanic/Latino | Anoxia/Asphyxiation |
| 2 | L5 | Female | 33 | White, non-Hispanic | Opioid overdose |
| 3 | L5 | Male | 55 | White; non-Hispanic | Anoxia/Seizure |

**Fig 5.**
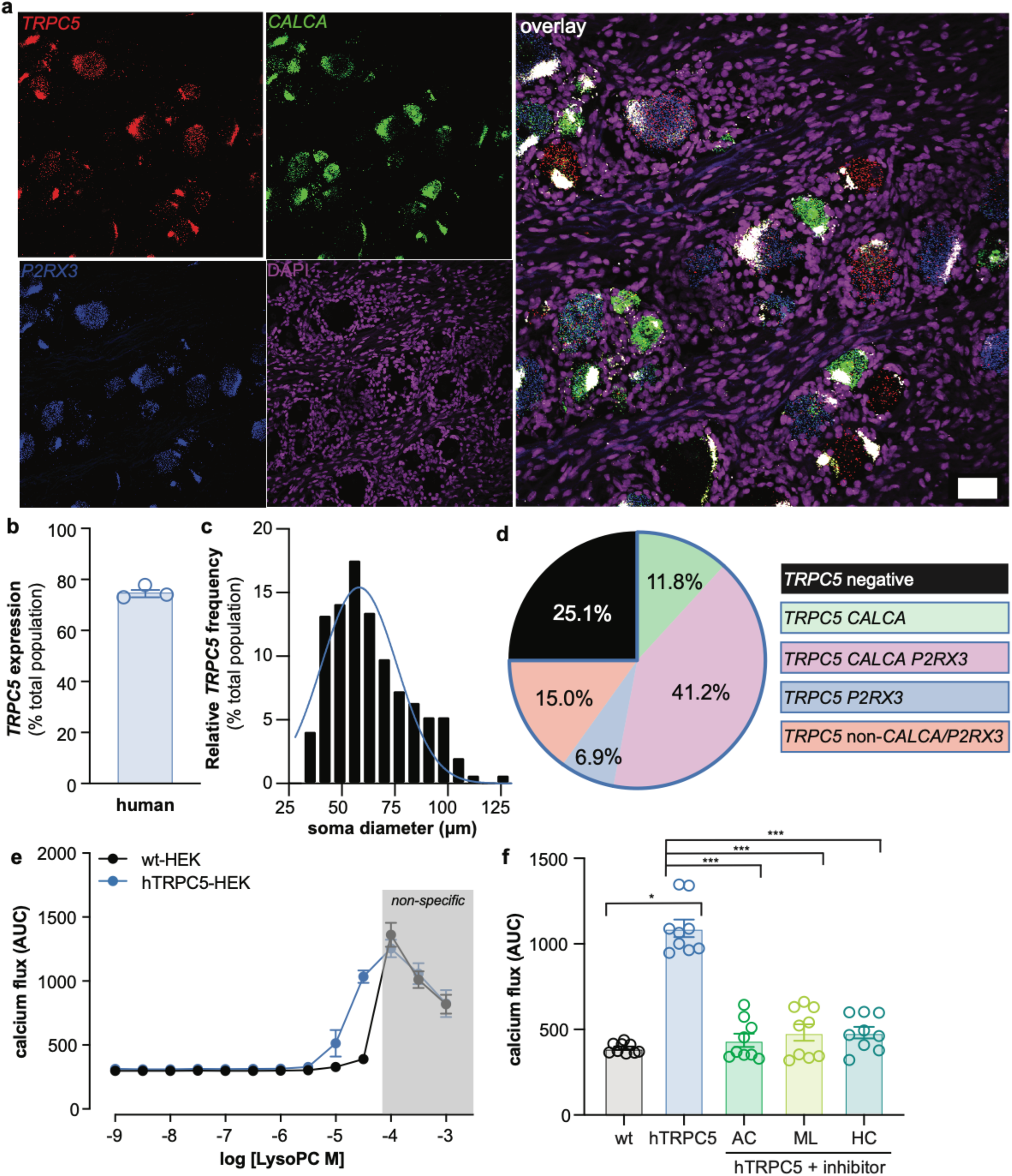
Human TRPC5 is expressed by the majority of DRG neurons and activated by LPC. **a**. Representative 20X image of human DRG neurons following RNAscope *in situ* hybridization for TRPC5 (red), CGRP (*CALCA*; green), P2XR3 (*P2RX3*; blue), and DAPI (magenta); scale bar: 50 µM. **b**. Percentage of DRG neurons that express TRPC5; n=3 humans. **c**. Frequency of TRPC5 positive neurons across population of human DRG neurons; n=438 neurons from 3 humans; bars are averages. **d**. Co-expression characterization of TRPC5-positive and TRPC5-negative neurons; pie wedges are averages; n=580 neurons from 3 humans. **e**. LPC-mediated calcium flux in HEK cells un-transfected or stably expressing hTRPC5; n=3 independent replicates performed in triplicate. **f**. Quantification of LPC (32 µM)-induced calcium flux in HEK cells un-transfected or stably expressing hTRPC5, and incubated with ML204 (320 µM), HC-070 (100 µM), or AC 1903 (320 µM); n=3 independent replicates performed in triplicate; Bonferroni post-hoc comparisons \**P*<0.05, \*\*\**P*<0.001. All data are mean ± SEM unless otherwise stated.

## Discussion

In this manuscript, we identified TRPC5 as a therapeutic target for persistent tactile and spontaneous pain. Using six different preclinical models, we were able to assess how TRPC5 contributes to pain arising from various pathophysiological sources. Peripheral administration of the potent TRPC5 inhibitors AC1903 or HC-070 reversed the established mechanical hypersensitivity associated with intraplantar CFA injections, hindpaw incision, and SCD. Similarly, global TPRC5 deletion prevented the development of mechanical allodynia and hyperalgesia associated with CFA injections, hindpaw incision, and paclitaxel treatment. Importantly, and unlike other mediators of mechanical allodynia (e.g. Piezo2, TRPA1, TRPV1, TRPV4), TRPC5 inhibition or genetic knockdown had no effect on naïve light touch or pain behaviors in uninjured mice, making TRPC5 a more promising druggable target than these other channels.

Since intraplantar administration of TRPC5 inhibitors was sufficient to alleviate tactile pain, we hypothesized that the anti-allodynic effects of TRPC5 manipulation were mediated by channels expressed in primary sensory afferents. Results from *ex vivo* and *in vitro* electrophysiological recordings supported this hypothesis. Following CFA injection, C fiber nociceptors from C5KO mice were less sensitive to mechanical stimulation than C fibers isolated from wildtype controls. Similarly, isolated DRG soma from SCD mice were less sensitive to mechanical stimulation when incubating with a TRPC5 inhibitor. But what was sensitizing TRPC5 across these different pain models? Was there a convergent mediator? To answer this question, we performed lipid mass spectroscopy on tissues harvested from CFA injected, hindpaw incised, nerve injured, and SCD mice. Tissue-specific increases in LPC, a byproduct of phosphatidylcholine metabolism *(37)*, were measured in all of the pain models that could be alleviated with TRPC5 inhibitors, therefore suggesting that LPC is a common endogenous mediator of TRPC5 activity and a biomarker for TRPC5-driven pain.

Despite not being involved in naïve neuronal mechanical sensitivity, TRPC5 does appear to mediate LPC-induced hypersensitivity. Local increases in LPC, a single fatty-acid chain phospholipid, likely sensitized TRPC5 to mechanical stimulation by altering neuronal membrane composition and resulting tension on membrane-embedded proteins. *In vitro* application of LPC increased the amplitude of mechanically-induced currents in wildtype TG neurons, but had no effect on the mechanical sensitivity of C5KO TG neurons. Similarly, dural application of LPC induced facial mechanical allodynia in wildtype mice but had no effect on C5KO controls. But do TRPC5 inhibitors retain their efficacy once tissue LPC levels normalize after injury? The previously described *in vitro* SCD patch clamping results would suggest that they do; > 24 hr after SCD DRG neurons were isolated from LPC-rich tissues, the mechanical hypersensitivity of these neurons was still reversed with TRPC5 inhibitors. Similarly, when isolated 48 hr after dural application of LPC, wildtype TG neurons retained lipid-induced mechanical hypersensitivity while C5KO neurons did not. This latent mechanical sensitization may result from increased trafficking of TRPC5 to the plasma membrane and subsequent activation by mechanical stimuli, G protein signaling, or endogenous agonists like protons *(16, 38)*. This same mechanism may be at play in many human pain conditions that are characterized by elevated LPC including migraine *(30)*. TRPC5 could be secondarily activated by the excessive G-protein coupled peptidergic signaling (e.g. CGRP, PACAP) *(39)* that occurs in migraine patients, thus making the channel a convergent target for migraine treatment.

In addition to decreasing mechanical hypersensitivity, TRPC5 inhibitors should also be considered for spontaneous pain management. Herein, application of exogenous LPC induced a conditioned place aversion in wildtype mice, but had no effect on C5KO mice. When sufficiently elevated, our data suggest that LPC can activate TRPC5 to induce spontaneous pain. CryoEM structures of mouse and human TRPC5 homomers have identified conserved phospholipid binding sites at the interface between individual monomer subunits *(31, 40, 41)*. Since AC1903, HC-070, and ML204, decreased LPC-induced TRPC5 activity in our heterologous expression system, these compounds may be effective therapies for LPC-related spontaneous pain *in vivo*. To extend the use of TRPC5 inhibitors for even more indications, additional endogenous TRPC5 activators should be comprehensively characterized.

Following the successful application of TRPC5 inhibitors in our preclinical models, we assessed TRPC5’s translational potential by using human tissues and expressing the human channel in heterologous systems. We observed uniform TRPC5 expression in DRG tissue from three different donors; TRPC5 was expressed in 75% of soma, most of which are putative nociceptors. The expression profile of TRPC5 was nearly identical to that observed for TRPV1, a high priority clinical target. First generation TRPV1 inhibitors were deemed unsuccessful when patients developed hyperthermia *(42)*, but second generation compounds effectively decreased osteoarthritis pain and are now in phase III clinical trials *(43)*. Since the expression profile of TRPC5 is just as wide as TRPV1, it is likely that compounds targeting these channels may be equally efficacious if administered for appropriate indications. In addition to widespread expression, TRPC5 target validity was further strengthened by the observation that hTRPC5 was activated by LPC *in vitro*. Like mTRPC5, LPC-induced hTRPC5 activity could be blocked with all of the TRPC5 inhibitors tested. Based on these *in vitro* data, TRPC5 inhibitors may be effective pain therapies in all of the following conditions that are associated with elevated LPC: rheumatoid arthritis *(44)*, osteoarthritis *(45)*, SCD *(34)*, fibromyalgia *(46)*, diabetes *(47)*, multiple sclerosis *(44)*, primary dysmenorrhea *(48)*, and lumbar spinal stenosis *(49)*. TRPC5 inhibitors are currently in phase I clinical trials for use in progressive kidney disease and anxiety. The anxiolytic effects of these compounds are proposedly mediated by TRPC5 expression in the amygdala *(50)*. In amygdala neurons, TRPC5 is primarily activated by GPCR signaling. The same GPCRs that activate TRPC5, namely group I metabotropic glutamate receptors *(50)*, are also involved in the top-down induction of mechanical allodynia *(51)* and visceral pain *(52)*. Therefore, when administered systemically, TRPC5 inhibitors may alleviate pain to an even greater extent by targeting both central and peripheral nervous system TRPC5 channels.

## Materials and Methods

### Animals

TRPC5 knockout *(53)* (C5KO), 129;S1 (C5KO wildtype), C57BL/6, Berkley SS*(54)* (SCD) and B6;129 (SCD wildtype) mice were bred in house and maintained on a 14:10 light/dark schedule with *ad libitum* access to rodent chow and water. Equally sized cohorts of male and female mice aged 8-26 weeks were used; 6-8 week old animals were used for dural injections. All protocols were in accordance with National Institutes of Health guidelines and were approved by the Institutional Animal Care and Use Committee at the Medical College of Wisconsin (Milwaukee, WI; protocol #383).

### Injury models

#### Complete Freud’s Adjuvant (CFA) injections

30 µL of stock CFA *(8, 55)* or sterile PBS was injected into the plantar aspect of one hind paw.

#### Hindpaw plantar incision

Postoperative pain was modeled using the hindpaw plantar incision procedure *(23, 56)*. Animals were maintained on inhalable isoflurane (1.5%) for the duration of the procedure (< 10 min). Starting 2 mm from the proximal edge of the heel, a 5 mm longitudinal incision was made through the glabrous skin. The underlying plantaris muscle was exposed, raised to rest on the tips of curved forceps, then incised longitudinally along the entire exposed length. The muscle was then repositioned, two sutures were used to close the incised skin, and bacitracin was applied to incision. Sham animals were anesthetized and had bacitracin applied to one hindpaw.

#### Spared tibial nerve injury (SNI)

A modified version of the spared nerve injury model *(57)* was used to model neuropathic pain. Animals were maintained on inhalable isoflurane (1.5%) for the duration of the procedure (<15 min). After incising the lateral skin of the thigh, the biceps femoris muscle was manipulated to expose the three distal branches of the sciatic nerve. The sural and common peroneal branches were ligated just distal to the branch point and transected 2-4 mm distal to the ligation; the tibial nerve was not ligated or cut. The overlying muscles were repositioned, and the skin incision was closed with staples. In sham animals, all branches of the sciatic nerve were exposed but not ligated or cut.

#### Paclitaxel treatment

Paclitaxel (Taxol) was dissolved in vehicle (16% ethanol, 16% Cremophor, 68% saline) to achieve a working solution of 0.8 mg/mL. Paclitaxel (8 mg/kg) or vehicle was injected intraperitoneally every other day over eight days.

#### Dural injections

Dural injections were performed as previously described *(28)*. Animals (≤ 8 weeks of age) were maintained on inhalable isoflurane (1.5%) for the duration of the procedure (<5 min). A 0.5 mm cannula was inserted through the soft tissue at the junction of the sagittal and lambdoid sutures; 5 µL of LPC, LPC+ AC1903, or pH 7.0 saline was infused through the cannula.

#### Behavioral tests

For all reflexive behavioral tests, animals were habituated to testing chamber and experimenter presence for >1 hr *(58)*. Experimenter was blinded to genotype, drug treatment, and/or injury model where applicable. Animals were randomly assigned to treatment groups.

#### von Frey mechanical allodynia testing

Hindpaw sensitivity to punctate mechanical stimulation was assessed via calibrated von Frey filaments (0.20-13.73 mN) and the up-down assessment method as previously described *(59, 60)*. The 50% withdrawal threshold was calculated for each paw as previously described *(61)*. Toe flaring without paw withdrawal was not considered a response. Calibrated von Frey filaments were also used to assess facial sensitivity to punctate mechanical stimulation as previously described *(28)*. Animals habituated to testing chambers (acrylic cups, 6.5 cm diameter x 7.0 cm height) for 2 hr on each day of the 2 days prior to start of testing. The 50% withdrawal threshold was calculated using the same analysis employed for hindpaw measurements.

#### Noxious needle testing

Hindpaw sensitivity to noxious punctate mechanical probing was assessed via needle stimulation as previously described *(62)*. A 25 Ga needle was pressed into the center of each hind paw with enough force to indent but not puncture the skin. Each paw was stimulated 10 times and the response frequency and characterization were recorded. Responses were characterized as null (no paw withdrawal), innocuous (simple withdrawal) or noxious (withdrawal accompanied by biting, licking, flicking, or additional attending to the paw).

#### Paintbrush testing

Hindpaw sensitivity to dynamic light touch was assessed via paintbrush stimulation as previously described *(63)*. A fine horsehair paintbrush was swept across the plantar surface of each hind paw from heel to toe pads, keeping the speed and force approximately constant between applications. Each paw was stimulated 10 times and the response frequency and characterization were reported as described in needle testing.

#### Conditioned place aversion

A 5- or 3-day protocol was used to assess the non-reflexive perception of LPC treatment. On preconditioning day, mice were placed into a two-chamber box. Individual chambers of the box could be visually differentiated by wall coverings (chamber 1: white polka dots on black background, chamber 2: equally sized black and white stripes on 30° diagonal). Chambers were connected via a retractable door. Each box was housed in a custom-built closed chamber fitted with a small circulating fan, white LED light strips, and overhead USB camera. Mice were allowed to freely move between the two chambers for 15 min. Animal movements were recorded by the overhead cameras and the total time spent in each chamber was analyzed with ANY-maze software. Animals spending <20% or >80% of the total trial in one of the two chambers were removed from the experiment on grounds of initial chamber bias *(64)*. Conditioning was completed in a biased fashion *(65, 66)*. In morning training sessions, all animals received a subcutaneous vehicle injection (1X PBS, pH 7.4; same volume as LPC injection), and were then immediately placed into their more preferred chamber from day 1 for 30 min. Animal movement was restricted to one chamber via the retractable door. In afternoon training sessions, all animals received subcutaneous injections of 1 or 5 µM LPC and were then immediately placed into their less preferred chamber from day 1 for 30 min. On postconditioning day 5, mice were again placed into the box and allowed to freely move between the two chambers for 15 min. Total time in each chamber on day 5 was recorded and differences between pre- and postconditioning times in each chamber were analyzed.

#### Paw edema measurements

Animals were anesthetized with inhalable isoflurane (1.5%) and the diameter of the tallest part of the paw was measured with digital calipers.

### Electrophysiology

#### Ex vivo teased tibial nerve recordings

Teased fiber tibial nerve recordings were completed as previously described *(67, 68)*. Seven days following CFA or vehicle injection, animals were anesthetized then sacrificed via cervical dislocation. The glabrous skin and tibial nerve from one leg was dissected and placed in a bath containing heated (32 ± 0.5°C), oxygenated buffer (pH of 7.45 ± 0.05) consisting of (in mM): 123 NaCl, 3.5 KCl, 2.0 CaCl_2_, 0.7 MgSO_4_, 1.7 NaH_2_PO_4_, 5.5 glucose, 7.5 sucrose 9.5 sodium gluconate and 10 HEPES. The nerve was placed into an adjacent chamber of the bath filled with mineral oil then teased into small bundles which were individually placed on silver wire electrode. Mechanically responsive receptive fields were identified by probing the corium with a blunt glass rod. For these experiments, activity was only recorded in mechanically responsive C fibers (conduction velocity <1.2 m/s *(69)*). Thresholds for mechanically-induced action potential firing were determined using a custom-built, feedback-controlled mechanical stimulator that applied a continuous force ramp from 0 to 100 mN. Firing frequencies were recorded during static force applications (2, 5, 10, 20, 40, 100, 150, and 200 mN for 10 s; 1 min inter-force interval). All data was recorded and analyzed in LabChart software. Experimenter was blinded to genotype until data analysis was complete.

#### Dorsal root ganglia (DRG) and trigeminal ganglia (TG) neuronal cultures

Mice were euthanized and sensory neurons were isolated from bilateral lumbar 1–6 ganglia or the bilateral trigeminal ganglia; culturing procedures were identical for both tissues. Ganglia were incubated with 1 mg/mL collagenase type IV for 40 min at 37°C and 5% CO_2_, then 0.05% trypsin for 45 min. Ganglia were mechanically dissociated and plated onto laminin-coated glass coverslips. Two hr after plating, neurons were fed with Dulbecco’s modified Eagle’s medium/Ham’s F12 medium supplemented with 10% heat-inactivated horse serum, 2 mM L-glutamine, 1% glucose, 100 units/ml penicillin, and 100 µg/ml streptomycin. Neurons grew overnight at 37°C, 5% CO_2_.

#### Whole-cell patch clamping

Coverslips were placed onto a Nikon Eclipse TE200 inverted microscope. Cells were continuously superfused with room temperature extracellular normal HEPES solution (pH 7.4 ± 0.05, and 310 ± 3 mOsm) containing (in mM): 140 NaCl, KCl, 2 CaCl_2_,1 MgCl_2_, 10 HEPES, and 10 glucose. Neurons (holding voltage −70 mV) were patched in voltage clamp mode with borosilicate glass pipettes filled with intracellular normal HEPES solution (pH 7.20 ± 0.05, and 290 ± 3 mOsm) containing (in mM): 135 KCl, 10 NaCl, 1 MgCl_2_, 1 EGTA, 0.2 NaGTP, 2.5 ATPNa_2_, and 10 HEPES. Cell capacitance and series resistance were kept below 10 MΩ for all cell types. Cell membranes were mechanically displaced (1.7 of 1.7 μm/V for 200 ms; 3 min inter-trial interval) by a second borosilicate glass pipette driven by a piezo stack actuator (pipette velocity: 106.25 μm/ms). All data was recorded in Pulse software. Current amplitude and type were analyzed in Fit Master software as previously described *(55)*. The mechanical threshold was designated as the first stimulation that elicited >20 pA inward current; mechanically insensitive cells were those that never exhibited an inward current >20 pA during mechanical stimulation. Cells were only included if the leak current stayed below 200 pA for ≥ three mechanical stimulations. For *in vitro* LPC or AC1903 experiments, neurons were exposed to chemical for a minimum of 5 min and maximum of 1.5 hr. Experimenter was blinded to genotype and treatment until data analysis was complete.

### Lipid mass spectroscopy

#### Tissue preparation

Hindpaw plantar skin was dissected 7 days following CFA injection or 5 days following incision. Sciatic nerve was isolated 10 days following SNI. Tissues were immediately frozen on dry ice after isolation. Serum was collected from all injury models at the same time points as follows: blood was obtained via cardiac puncture from deeply anesthetized mice. Blood was centrifuged at 1500 rpm, 4°C, for 15 min. Serum was pipetted into sterile tube then immediately frozen on dry ice. Samples were then shipped to the Northwest Metabolomics Research Center for further processing.

#### Sample preparation

Lipids were extracted from samples as previously described *(70)*. Samples were reconstituted in 250 µL Lipidyzer running buffer composed of dichloromethane:methanol with 10 mM ammonium acetate.

#### Mass spectrometry

Quantitative lipidomics was performed with the Sciex Lipidyzer platform consisting of Shimadzu Nexera X2 LC-30AD pumps, a Shimadzu Nexera X2 SIL-30AC autosampler, and a Sciex QTRAP® 5500 mass spectrometer equipped with SelexION® for differential mobility spectrometry (DMS). 1-propanol was used as the chemical modifier for the DMS. Samples were introduced to the mass spectrometer by flow injection analysis at 8 µL/min. Lipids were analyzed in negative ESI mode using multiple reaction monitoring (MRM). A total of 26 LPC lipids were targeted in the analysis.

#### Data processing

Data was acquired and processed using Analyst 1.6.3 and Lipidomics Workflow Manager 1.0.5.0. Stucky Lab members analyzed the data in a blinded fashion.

#### TRPC5 Stable Cell Line Generation and Fluo-4 Calcium Flux Assays

Stable cell lines were generated in HEK293 (ATCC, mycoplasma free) using a pCMV vector expressing either 1µg of mouse or human TRPC5 (Origene) co-transfected with 7µg of pBabe Puro vector for rapid stable selection. Cell lines were selected and maintained DMEM containing 10% FBS, 5 µg/mL Puromycin (GoldBio), and 750 µg/mL G418 (GoldBio). On the day before assay, cells were seeded into 384-well poly-L-lysine-coated black plates at a density of 10,000 cells/well in DMEM containing 1% dialyzed FBS. On the day of the assay, media was decanted, and the cells were incubated with Fluo-4 Direct dye (Invitrogen, 20 µl/well) for 1 h at 37 °C, which was reconstituted in drug buffer in a 1:10 dilution (20 mM HEPES-buffered HBSS, pH 7.4) containing 2.5 mM probenecid. After dye load, cells were allowed to equilibrate to room temperature for 15 minutes, and then placed in a FLIPR^TETRA^ fluorescence imaging plate reader (Molecular Dynamics). Ligand dilutions were prepared at 5X final concentration in drug buffer (20 mM HEPES-buffered HBSS, pH 7.4) containing 0.1% BSA and 0.01% ascorbic acid, final concentration. Ligand dilutions were aliquoted into 384-well plastic plates and placed in the FLIPR^TETRA^ for drug stimulation. For competition antagonist assays, plates were challenged with either LPC (32-60 µM) or Englerin A (100 nM) and prepared at 6X final concentration in drug buffer (20 mM HEPES-buffered HBSS, pH 7.4) containing 0.1% BSA and 0.01% ascorbic acid, final concentration. Fluorescence for the FLIPR^TETRA^ were programmed to read baseline fluorescence for 10 s (1 read/s), and afterward 5 µl of drug per well was added and read for a total of 5 min (1 read/s). Fluorescence in each well was normalized to the average of the first 10 reads for baseline fluorescence, and then the area under the curve (AUC) was determined and calculated. AUC was plotted as a function of ligand concentration, and data were analyzed as non-linear regression using “log(agonist) vs. response—variable slope” in GraphPad Prism 5.0 to yield Emax and EC_50_ parameter estimates.

#### Chemical matter

AC1903 (provided by Dr. Corey Hopkins *(18)*; 2.5 µg/µL) was gently heated in 10% EtOH, 40% PEG400, 50% saline vehicle until dissolved. Lysophosphatidylcholine (LPC) was purchased from Sigma Aldrich and dissolved in 0.02% EtOH vehicle. ML 204 was purchased from Sigma Aldrich and dissolved in 12.5% DMSO vehicle. HC-070 was purchased from MedChemExpress and dissolved in 0.125% DMSO vehicle *(32)*. (-)-Englerin A was purchased from Sigma Aldrich*(71)*. For in vitro experiments, all compounds were dissolved in DMSO at 100 mM, except LPC was dissolved in ethanol.

### RNAscope

#### Tissue preparation

All human tissue procurement procedures were approved by the Institutional Review Boards at the University of Texas at Dallas. Human dorsal root ganglion (L5) were collected, frozen on dry ice and stored in a -80°C freezer. Donor information is provided in **Table 4**. The human DRGs were gradually embedded with OCT in a cryomold by adding small volumes of OCT over dry ice to avoid thawing. All tissues were cryostat sectioned at 20 μm onto SuperFrost Plus charged slides. Sections were only briefly thawed in order to adhere to the slide but were immediately returned to the -20°C cryostat chamber until completion of sectioning. The slides were then immediately utilized for histology.

#### RNAscope *in situ* hybridization

RNAscope *in situ* hybridization multiplex version 1 was performed as instructed by Advanced Cell Diagnostics (ACD). Slides were removed from the cryostat and immediately transferred to cold (4°C) 10% formalin for 15 minutes. The tissues were then dehydrated in 50% ethanol (5 min), 70% ethanol (5 min) and 100% ethanol (10 min) at room temperature. The slides were air dried briefly and then boundaries were drawn around each section using a hydrophobic pen (ImmEdge PAP pen; Vector Labs). When hydrophobic boundaries had dried, protease IV reagent was added to each section until fully covered and incubated for 2-5 minutes at room temperature. The protease IV incubation period was optimized for the specific lot of Protease IV reagent and for each DRG as recommended by ACD. Slides were washed briefly in 1X phosphate buffered saline (PBS, pH 7.4) at room temperature. Each slide was then placed in a prewarmed humidity control tray (ACD) containing dampened filter paper and a 50:1:1 dilution (as directed by ACD due to stock concentrations) of *TRPC5* (ACD Cat # 427651; Channel 1), *CALCA* (ACD Cat # 605551; Channel 2), *P2RX3* (ACD Cat # 406301; Channel 3) was pipetted onto each section until fully submerged. This was performed one slide at a time to avoid liquid evaporation and section drying. The humidity control tray was placed in a HybEZ oven (ACD) for 2 hours at 40°C. Following probe incubation, the slides were washed two times in 1X RNAscope wash buffer and returned to the oven for 30 minutes after submersion in AMP-1 reagent. Washes and amplification were repeated using AMP-2, AMP-3 and AMP-4 reagents with a 15-min, 30-min, and 15-min incubation period, respectively. AMP-4 ALT C (Channel 1 = Atto 550, Channel 2 = Atto 647, Channel 3 = Alexa 488) was used for all experiments. Slides were then washed two times in 0.1M phosphate buffer (PB, pH7.4). Human slides were incubated in DAPI (1/5000) in 0.1M PB for 1 min before being washed, air dried, and cover-slipped with Prolong Gold Antifade mounting medium.

#### Tissue quality check

All human DRGs were checked for RNA quality by using a positive control probe cocktail (ACD) which contains probes for high, medium and low-expressing mRNAs that are present in all cells (ubiquitin C > Peptidyl-prolyl cis-trans isomerase B > DNA-directed RNA polymerase II subunit RPB1). DRGs that showed signal for all 3 positive control probes were used to generate experimental data. A negative control probe against the bacterial DapB gene (ACD) was used to check for non-specific/background label.

#### Image Analysis

DRG sections were imaged on an Olympus FV3000 confocal microscope at 20X magnification. 3-4 20X images were acquired of each human DRG section, and 3 sections were imaged per human donor. The acquisition parameters were set based on guidelines for the FV3000 provided by Olympus. In particular, the gain was kept at the default setting 1, HV ≤ 600, offset = 4, and laser power ≤ 15%. The raw image files were brightened and contrasted in Olympus CellSens software (v1.18), and then analyzed manually one cell at a time for expression of each gene target. Cell diameters were measured using the polyline tool. Total neuron counts for human samples were acquired by counting all of the probe-labeled neurons and all neurons that were clearly outlined by DAPI (satellite cell) signal and contained lipofuscin in the overlay image.

Large globular structures and/or signal that auto-fluoresced in all 3 channels (488, 550, and 647; appears white in the overlay images) was considered to be background lipofuscin and was not analyzed. Aside from adjusting brightness/contrast, we performed no digital image processing to subtract background. We attempted to optimize automated imaging analysis tools for our purposes, but these tools were designed to work with fresh, low background rodent tissues, not human samples taken from older organ donors. As such, we chose to implement a manual approach in our imaging analysis in which we used our own judgement of the negative/positive controls and target images to assess mRNA label. Images were not analyzed in a blinded fashion.

## Data analysis

Individual data points were presented whenever possible in addition to group means ± SEM. Data were analyzed using GraphPad Prism 8; results were considered statistically significant when *P*<0.05. For behavioral experiments, data from each sex were independently analyzed. No significant differences were noted so data were re-analyzed with sexes combined.

## Acknowledgements

We thank Dr. Corey Hopkins for generously providing AC1903, the laboratory of Dr. Ellen Lumpkin for technical advice, Dr. David Clapham for comments on the manuscript, the Neuroscience Research Center at the Medical College of Wisconsin for access to behavioral equipment, and the Northwest Metabolomics Research Center for lipid mass spectroscopy.

## Author contributions

KES planned, performed, and analyzed behavioral and electrophysiology experiments, wrote and edited the manuscript. FM planned, performed, and analyzed behavioral and electrophysiology experiments and edited the manuscript. SIS planned, performed, and analyzed RNAscope experiment, wrote and edited the manuscript. LJL planned, performed, and analyzed in vitro experiments, and edited the manuscript. ARM planned, performed, and analyzed behavioral experiments and edited the manuscript. ZRP planned, performed, and analyzed in vitro experiments, and edited the manuscript. CMM performed behavioral experiments and edited the manuscript. GD assisted in experimental planning and manuscript editing. TJP assisted in experimental planning and manuscript editing. JDM planned, performed, and analyzed in vitro experiments, and edited the manuscript. CLS assisted in experimental planning and manuscript editing.

## Data and materials availability

All data associated with this study are available in the main text or the supplementary materials.

---

**Fig S1.**
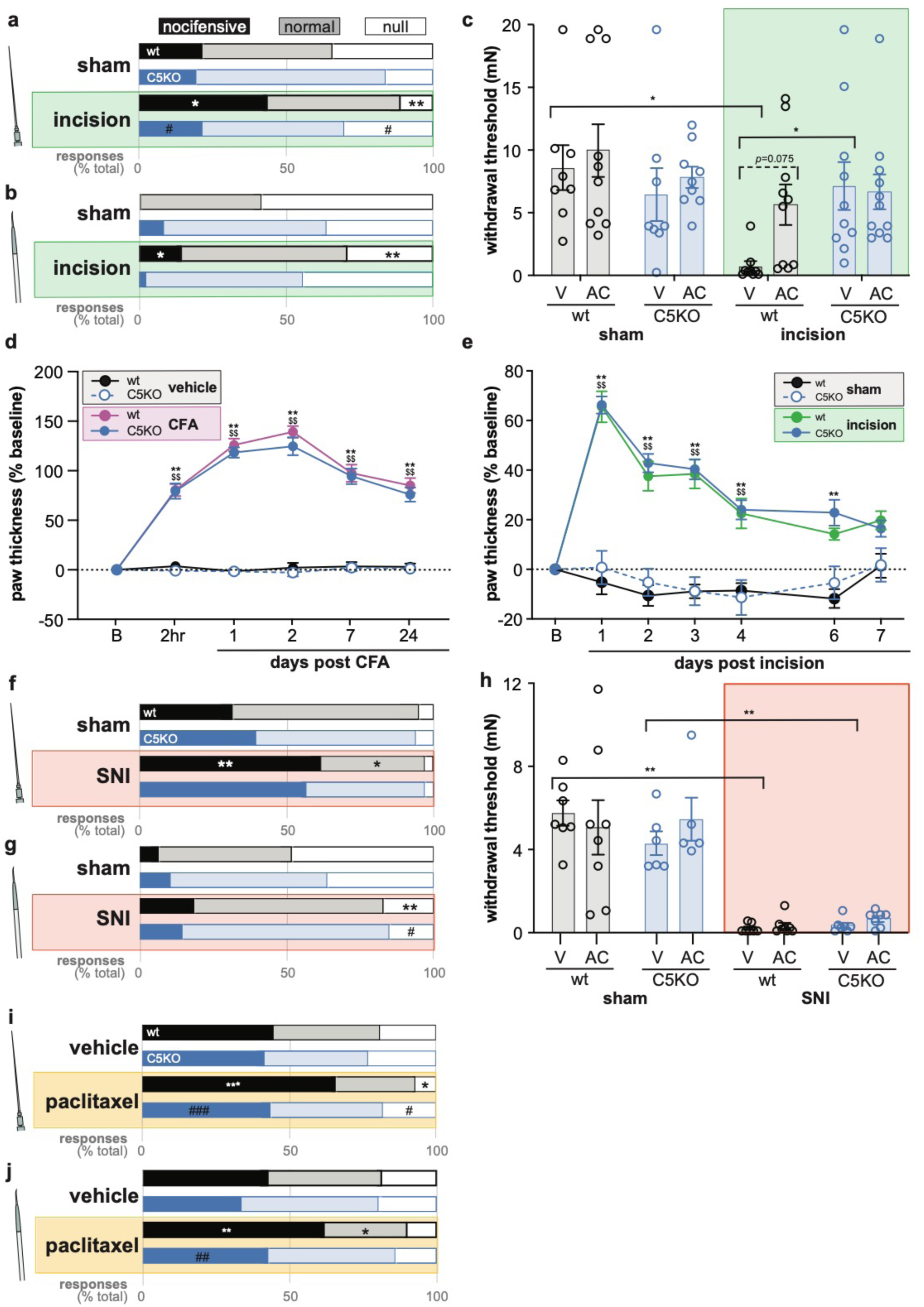
TRPC5 contributes to persistent mechanical hypersensitivity in select inflammatory and neuropathic pain models. **a**. Response characterization to noxious needle hindpaw stimulation 4 days post-hindpaw plantar incision; n=8-9 mice. **b**. Response characterization to paintbrush swiping across hindpaw 4 days post-hindpaw plantar incision; n=8-9 mice. **c**. Mechanical withdrawal thresholds on day 5 post-hindpaw plantar incision, 60 min following intraplantar injection of AC1903 (AC) or vehicle (V); n=8-10 mice; Bonferroni post-hoc comparison \**P*<0.05. **d**. Percent change from baseline (B) in paw thickness post-CFA injection; n=9-10 mice. **e**. Percent change from baseline in paw thickness post-hindpaw plantar incision; n=9-10 mice. **f**. Response characterization to noxious needle hindpaw stimulation 24 days post-SNI; n=7-10 mice. **g**. Response characterization to paintbrush swiping across hindpaw 24 days post-SNI; n=7-10 mice. **h**. Mechanical withdrawal thresholds on day 28 post-SNI, 60 min following intraplantar injection of AC1903 vehicle; n=5-8 mice; Bonferroni post-hoc comparison \*\**P*<0.01. **i**. Response characterization to noxious needle hindpaw stimulation 16 days post-start of paclitaxel treatment; n=8-10 mice. **j**. Response characterization to paintbrush swiping across hindpaw 16 days post-start of paclitaxel treatment; n=8-10 mice. Bars in **a**, b, f, **g, I**, and **j** are group averages. All other data are mean ± SEM. Bonferroni post-hoc comparisons for all panels unless otherwise stated.: \**P*<0.05, \*\**P*<0.01, \*\*\**P*<0.001 wildtype control vs. injury; ^$$^*P*<0.01, C5KO control vs. injury; ^#^*P*<0.05, ^##^*P*<0.01, ^###^*P*<0.001 injury C5KO vs. wildtype.

**Fig S2.**
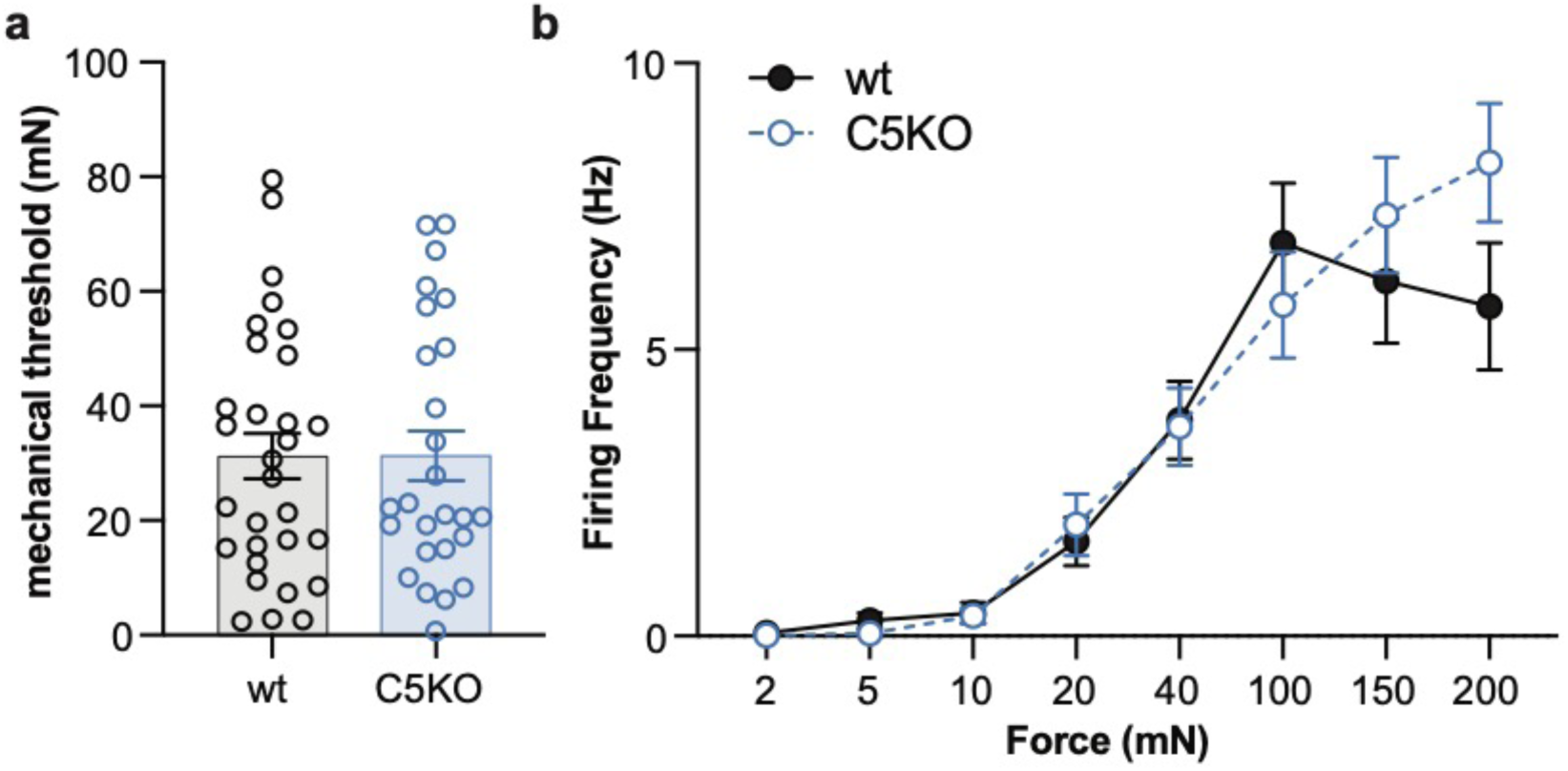
C fibers from TRPC5 knockout (C5KO) mice exhibit normal responses to receptive field mechanical stimulation. **a**. Mechanical thresholds of C fibers from vehicle-injected wildtype and C5KO mice; n=27-30 units from 5 mice. **b**. Mean mechanically-induced firing rates of C fibers from vehicle-injected wildtype and C5KO mice; n=27-30 units from 5 mice.

**Fig S3.**
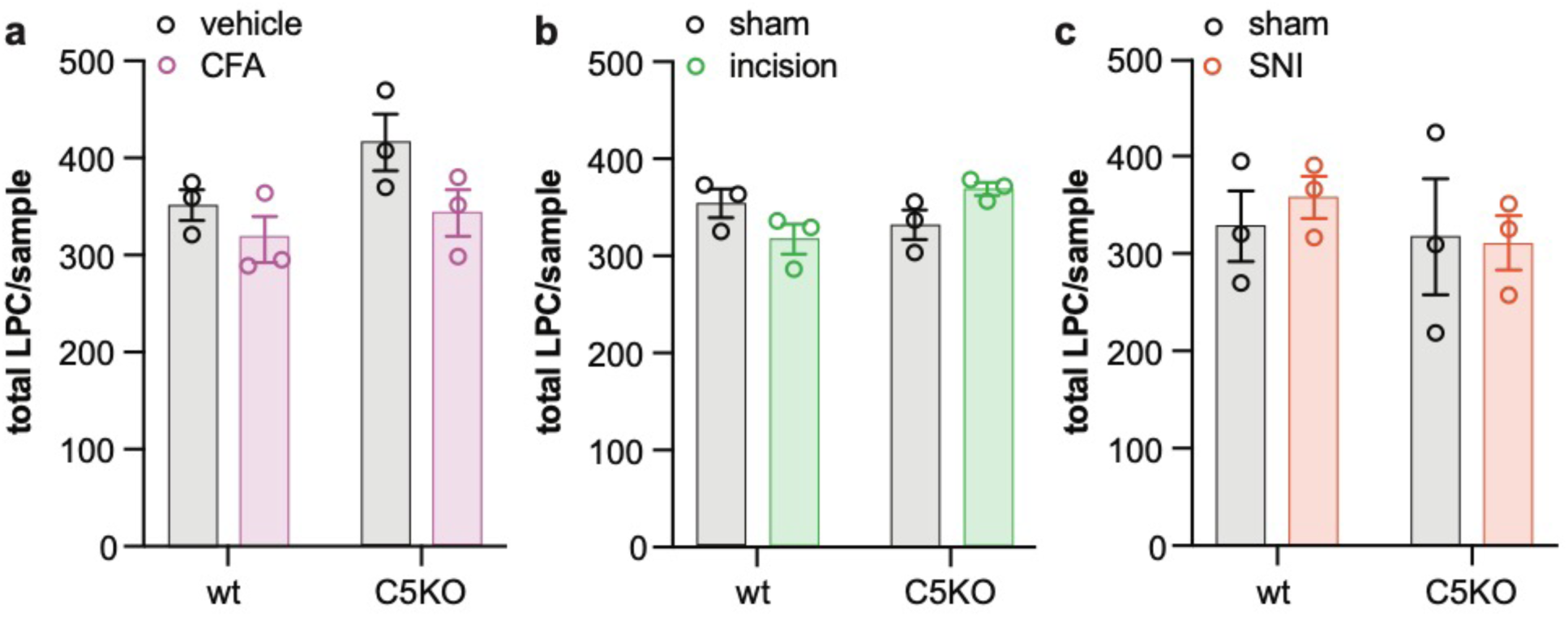
Serum LPC levels are unchanged following peripheral injury. **a**. Relative amount of LPC in serum 7 days post-CFA injection; n=3 mice. **b**. Relative amount of LPC in serum 5 days post-hindpaw plantar incision; n=3 mice. **c**. Relative amount of LPC in serum 10 days post-SNI; n=3 mice.

**Fig S4.**
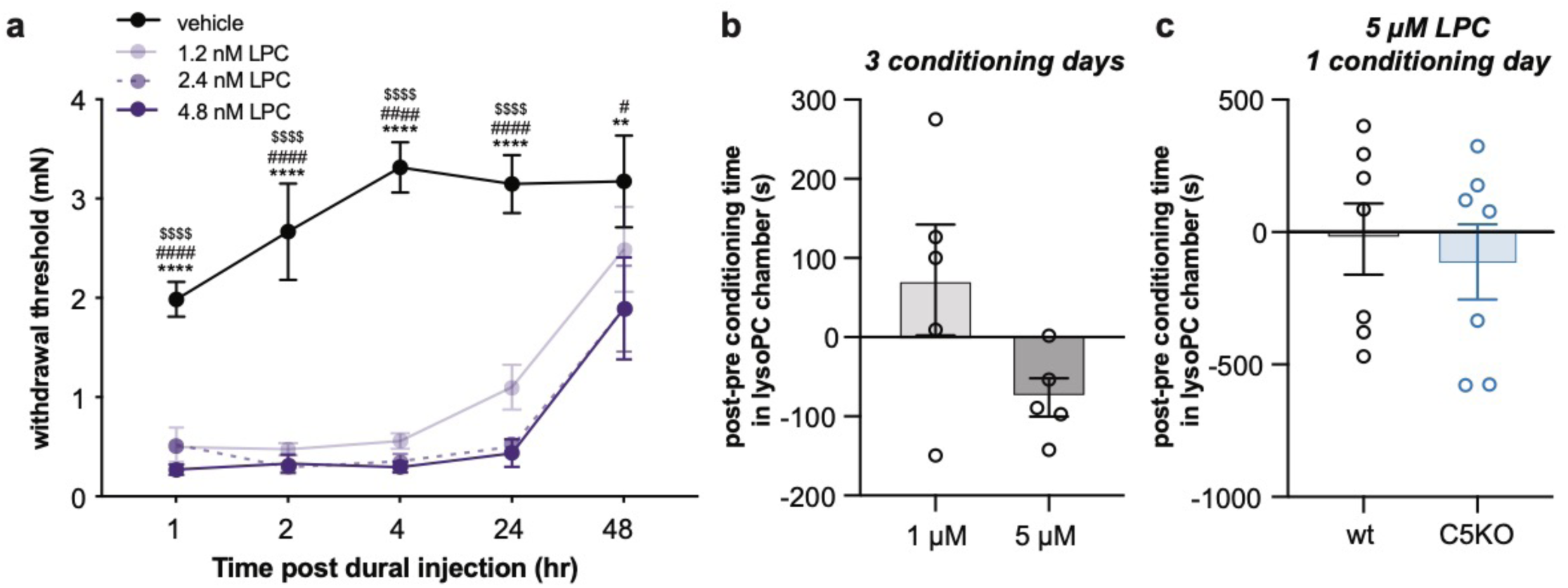
Exogenous application of LPC induces mechanical hypersensitivity and spontaneous pain. **a**. Periorbital mechanical withdrawal thresholds of wildtype mice following dural application of LPC; n=8-10 mice; Bonferroni post-hoc comparisons: \*\**P*<0.01, \*\*\*\**P*<0.0001 4.8 nM vs. vehicle; ^#^*P*<0.05, ^####^*P*<0.0001 2.4 nM vs. vehicle, ^$$$$^*P*<0.0001 1.2 nM vs vehicle. **b**. Difference in LPC chamber time between pre- and post-conditioning trials; n=5 mice, 3 days of conditioning. **c**. Difference in LPC chamber time between pre- and post-conditioning trials; n=7 mice, 1 day of conditioning, 5 µM LPC. All data are mean ± SEM.

**Fig S5.**
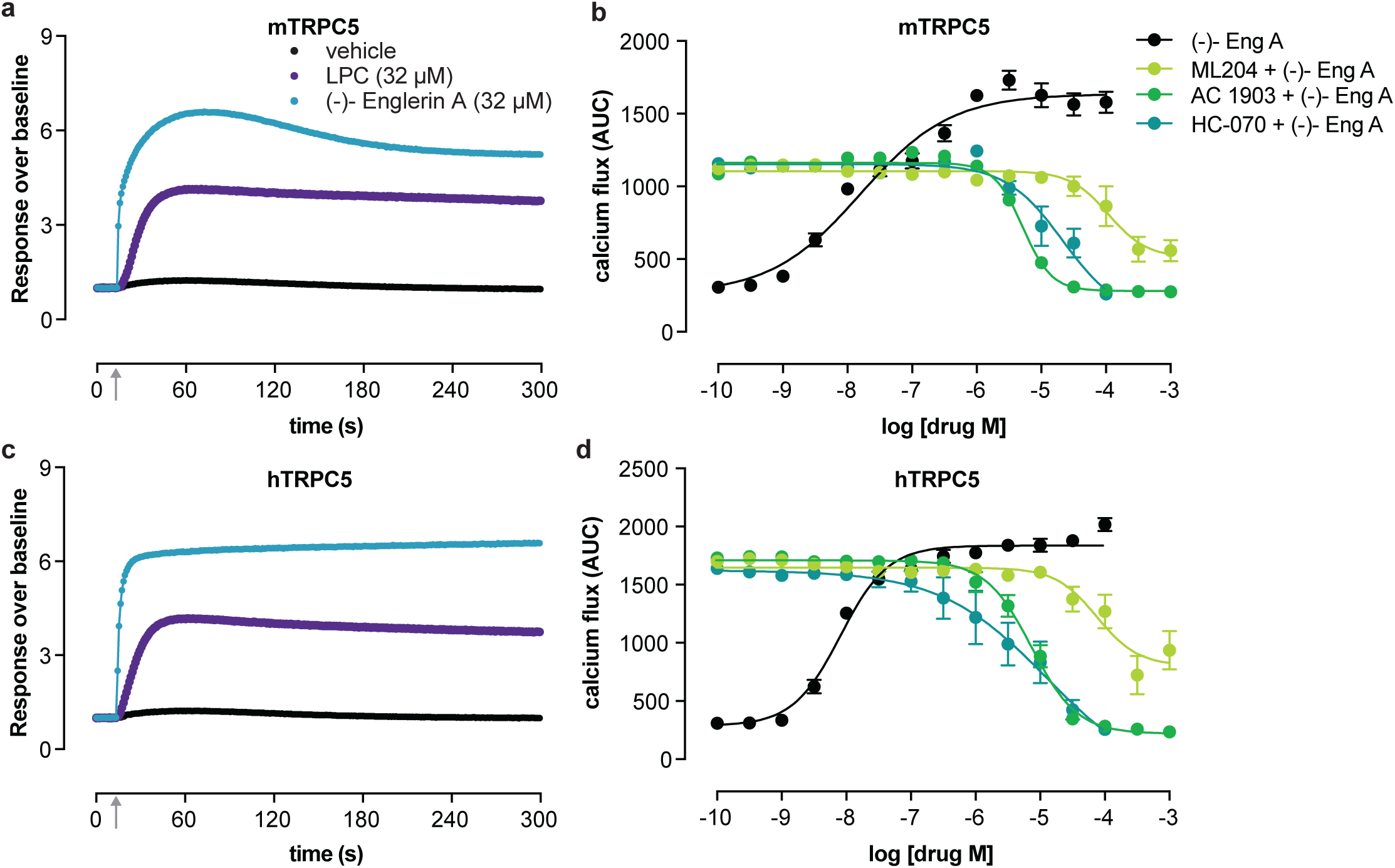
LPC and (-)-Englerin-A directly activate mouse and human TRPC5. **a**. Time course of calcium flux in HEK cells stably expressing mTRPC5 when exposed to LPC, (-)-Englerin A, or vehicle; gray arrow: chemical application (time = 10 s). **b**. mTRPC5 activation by 100 nM (-)-Eng A (black, EC_50_ = 16.0 nM) is blocked by ML204 (IC_50_ = 109 µM), HC-070 (IC_50_ = 22 µM), and AC 1903 (IC_50_ = 5.2 µM). **c**. Time course of calcium flux in HEK cells stably expressing hTRPC5 when exposed to LPC, (-)-Englerin A, or vehicle; gray arrow: chemical application (time = 10 s). **d**. hTRPC5 activation by 100 nM (-)-Eng A (black, EC_50_ = 8.2 nM) is blocked by ML204 (IC_50_ = 75 µM), HC-070 (IC_50_ = 12 µM), and AC 1903 (IC_50_ = 7.4 µM). Data represent mean and SEM from n=4-5 independent experiments performed in triplicate.

## References

1. S. Lolignier, N. Eijkelkamp, J. N. Wood, Mechanical allodynia *Pflugers Arch*. Eur. J. Physiol. 467, 133–139 (2014).

2. C. J. Woolf, Central sensitization: Implications for the diagnosis and treatment of pain Pain 152 (2011), doi: 10.1016/j.pain.2010.09.030.

3. S. R. Eid, E. D. Crown, E. L. Moore, H. A. Liang, K. C. Choong, S. Dima, D. A. Henze, S. A. Kane, M. O. Urban, HC-030031, a TRPA1 selective antagonist, attenuates inflammatory- and neuropathy-induced mechanical hypersensitivity, Mol. Pain 4 (2008), doi: 10.1186/1744-8069-4-48.

4. M. Szczot, J. Liljencrantz, N. Ghitani, A. Barik, R. Lam, J. H. Thompson, D. Bharucha-Goebel, D. Saade, A. Necaise, S. Donkervoort, A. R. Foley, T. Gordon, L. Case, M. C. Bushnell, C. G. Bönnemann, A. T. Chesler, PIEZO2 mediates injury-induced tactile pain in mice and humans, Sci. Transl. Med. (2018), doi: 10.1126/scitranslmed.aat9892.

5. S. E. Murthy, M. C. Loud, I. Daou, K. L. Marshall, F. Schwaller, J. Kühnemund, A. G. Francisco, W. T. Keenan, A. E. Dubin, G. R. Lewin, A. Patapoutian, The mechanosensitive ion channel Piezo2 mediates sensitivity to mechanical pain in mice, Sci. Transl. Med. (2018), doi: 10.1126/scitranslmed.aat9897.

6. K. Y. Kwan, A. J. Allchorne, M. A. Vollrath, A. P. Christensen, D. S. Zhang, C. J. Woolf, D. P. Corey, TRPA1 Contributes to Cold, Mechanical, and Chemical Nociception but Is Not Essential for Hair-Cell Transduction, Neuron 50, 277–289 (2006).

7. M. Petrus, A. M. Peier, M. Bandell, S. W. Hwang, T. Huynh, N. Olney, T. Jegla, A. Patapoutian, A role of TRPA1 in mechanical hyperalgesia is revealed by pharmacological inhibition, Mol. Pain (2007), doi: 10.1186/1744-8069-3-40.

8. R. C. Lennertz, E. A. Kossyreva, A. K. Smith, C. L. Stucky, TRPA1 mediates mechanical sensitization in nociceptors during inflammation., PLoS One 7, e43597 (2012).

9. J. P. Dunham, S. Kelly, L. F. Donaldson, Inflammation reduces mechanical thresholds in a population of transient receptor potential channel A1-expressing nociceptors in the rat, Eur. J. Neurosci. 27, 3151–3160 (2008).

10. B. Coste, J. Mathur, M. Schmidt, T. J. Earley, S. Ranade, M. J. Petrus, A. E. Dubin, A. Patapoutian, Piezo1 and Piezo2 are essential components of distinct mechanically activated cation channels, Science (80-.). 330, 55–60 (2010).

11. A. T. Chesler, M. Szczot, D. Bharucha-Goebel, M. Čeko, S. Donkervoort, C. Laubacher, L. H. Hayes, K. Alter, C. Zampieri, C. Stanley, A. M. Innes, J. K. Mah, C. M. Grosmann, N. Bradley, D. Nguyen, A. R. Foley, C. E. Le Pichon, C. G. Bönnemann, The Role of *PIEZO2* in Human Mechanosensation, N. Engl. J. Med. 375, 1355–1364 (2016).

12. P. C. Kerstein, D. del Camino, M. M. Moran, C. L. Stucky, Pharmacological blockade of TRPA1 inhibits mechanical firing in nociceptors, Mol. Pain 5 (2009), doi: 10.1186/1744-8069-5-19.

13. K. Y. Kwan, J. M. Glazer, D. P. Corey, F. L. Rice, C. L. Stucky, TRPA1 modulates mechanotransduction in cutaneous sensory neurons., J. Neurosci. 29, 4808–19 (2009).

14. K. J. Zappia, C. L. O’Hara, F. Moehring, K. Y. Kwan, C. L. Stucky, Sensory neuron-specific deletion of TRPA1 results in mechanical cutaneous sensory deficits, eneuro 4, ENEURO.0069-16.2017 (2017).

15. K. Zimmermann, J. K. Lennerz, A. Hein, A. S. Link, J. S. Kaczmarek, M. Delling, S. Uysal, J. D. Pfeifer, A. Riccio, D. E. Clapham, Transient receptor potential cation channel, subfamily C, member 5 (TRPC5) is a cold-transducer in the peripheral nervous system, Proc. Natl. Acad. Sci. 108, 18114–18119 (2011).

16. M. Semtner, M. Schaefer, O. Pinkenburg, T. D. Plant, Potentiation of TRPC5 by protons, J. Biol. Chem. 282, 33868–33878 (2007).

17. P. K. Flemming, A. M. Dedman, S. Z. Xu, J. Li, F. Zeng, J. Naylor, C. D. Benham, A. N. Bateson, K. Muraki, D. J. Beech, Sensing of lysophospholipids by TRPC5 calcium channel, J. Biol. Chem. 281, 4977–4982 (2006).

18. Y. Zhou, P. Castonguay, E. H. Sidhom, A. R. Clark, M. Dvela-Levitt, S. Kim, J. Sieber, N. Wieder, J. Y. Jung, S. Andreeva, J. Reichardt, F. Dubois, S. C. Hoffmann, J. M. Basgen, M. S. Montesinos, A. Weins, A. C. Johnson, E. S. Lander, M. R. Garrett, C. R. Hopkins, A. Greka, A small-molecule inhibitor of TRPC5 ion channels suppresses progressive kidney disease in animal models, Science (80-.). 358, 1332–1336 (2017).

19. T. J. Brennan, E. P. Vandermeulen, G. F. Gebhart, Characterization of a rat model of incisional pain, Pain 64, 493–502 (1996).

20. A. Gomis, S. Soriano, C. Belmonte, F. Viana, Hypoosmotic- and pressure-induced membrane stretch activate TRPC5 channels, J. Physiol. 586, 5633–5649 (2008).

21. B. Shen, C. O. Wong, O. C. Lau, T. Woo, S. Bai, Y. Huang, X. Yao, Plasma membrane mechanical stress activates TRPC5 channels, PLoS One 10, 1–18 (2015).

22. A. D. Weyer, K. J. Zappia, S. R. Garrison, C. L. O’Hara, A. K. Dodge, C. L. Stucky, Nociceptor Sensitization Depends on Age and Pain Chronicity., eNeuro 3 (2016), doi: 10.1523/ENEURO.0115-15.2015.

23. A. M. Cowie, A. D. Menzel, C. O′Hara, M. W. Lawlor, C. L. Stucky, NOD-like receptor protein 3 inflammasome drives postoperative mechanical pain in a sex-dependent manner, Pain 160, 1794–1816 (2019).

24. A. K. Smith, C. L. O’Hara, C. L. Stucky, Mechanical sensitization of cutaneous sensory fibers in the spared nerve injury mouse model, Mol. Pain 9, 1744-8069-9–61 (2013).

25. H. S. Shim, C. Bae, J. Wang, K. H. Lee, K. M. Hankerd, H. K. Kim, J. M. Chung, J. H. La, Peripheral and central oxidative stress in chemotherapy-induced neuropathic pain, Mol. Pain 15 (2019), doi: 10.1177/1744806919840098.

26. P. K. Flemming, A. M. Dedman, S. Z. Xu, J. Li, F. Zeng, J. Naylor, C. D. Benham, A. N. Bateson, K. Muraki, D. J. Beech, Sensing of lysophospholipids by TRPC5 calcium channel, J. Biol. Chem. 281, 4977–4982 (2006).

27. S.-H. Law, M.-L. Chan, G. K. Marathe, F. Parveen, C.-H. Chen, L.-Y. Ke, An Updated Review of Lysophosphatidylcholine Metabolism in Human Diseases, Int. J. Mol. Sci. 20, 1149 (2019).

28. C. C. Burgos-Vega, L. D. Quigley, G. Trevisan dos Santos, F. Yan, M. Asiedu, B. Jacobs, M. Motina, N. Safdar, H. Yousuf, A. Avona, T. J. Price, G. Dussor, Non-invasive dural stimulation in mice: A novel preclinical model of migraine Cephalalgia (2018), doi: 10.1177/0333102418779557.

29. R. Burstein, D. Yarnitsky, I. Goor-Aryeh, B. J. Ransil, Z. H. Bajwa, An association between migraine and cutaneous allodynia, Ann. Neurol. 47, 614–624 (2000).

30. C. Ren, J. Liu, J. Zhou, H. Liang, Y. Wang, Y. Sun, B. Ma, Y. Yin, Lipidomic analysis of serum samples from migraine patients, Lipids Health Dis. 17, 1–9 (2018).

31. K. Song, M. Wei, W. Guo, Y. Kang, J.-X. Wu, L. Chen, Structural basis for human TRPC5 channel inhibition by two distinct inhibitors, bioRxiv, 2020.04.21.052910 (2020).

32. S. Just, B. Chenard, A. Cecl, T. Strassmaier, J. Chong, N. Blair, R. Gallaschun, D. Camino, S. Cantin, M. D’Amours, C. Eickmeier, C. Fanger, C. Hecker, D. Hessler, B. Hengerer, K. Kroker, S. Malekiani, R. Mihalek, J. McLaughlin, G. Rast, W. J, A. Sauer, C. Pryce, M. Moran, Treatment with HC-070, a potent inhibitor of TRPC4 and TRPC5, leads to anxiolytic and antidepressant effects in mice (2018).

33. M. Miller, J. Shi, Y. Zhu, M. Kustov, J. Bin Tian, A. Stevens, M. Wu, J. Xu, S. Long, P. Yang, A. Zholos V, J. M. Salovich, C. D. Weaver, C. R. Hopkins, C. W. Lindsley, O. McManus, M. Li, M. X. Zhu, Identification of ML204, a novel potent antagonist that selectively modulates native TRPC4/C5 ion channels, J. Biol. Chem. 286, 33436–33446 (2011).

34. H. Wu, M. Bogdanov, Y. Zhang, K. Sun, S. Zhao, A. Song, R. Luo, N. F. Parchim, H. Liu, A. Huang, M. G. Adebiyi, J. Jin, D. C. Alexander, M. V Milburn, M. Idowu, H. S. Juneja, R. E. Kellems, W. Dowhan, Y. Xia, Hypoxia-mediated impaired erythrocyte Lands’ Cycle is pathogenic for sickle cell disease, Sci. Rep. 6, 29637 (2016).

35. J. S. Mogil, Animal models of pain: Progress and challenges Nat. Rev. Neurosci. 10, 283–294 (2009).

36. S. Shiers, R. M. Klein, T. J. Price, Quantitative differences in neuronal subpopulations between mouse and human dorsal root ganglia demonstrated with RNAscope in situ hybridization, Pain (2020), doi: 10.1097/j.pain.0000000000001973.

37. I. Kudo, M. Murakami, Phospholipase A2 enzymes, Prostaglandins Other Lipid Mediat. 68–69, 3–58 (2002).

38. D. J. Beech, Canonical transient receptor potential 5, Handb. Exp. Pharmacol. 179, 109–123 (2007).

39. E. A. Kaiser, A. F. Russo, CGRP and migraine: Could PACAP play a role too? Neuropeptides 47, 451–461 (2013).

40. D. J. Wright, K. J. Simmons, R. M. Johnson, D. J. Beech, S. P. Muench, R. S. Bon, Cryo-EM structures of human TRPC5 reveal interaction of a xanthine-based TRPC1/4/5 inhibitor with a conserved lipid binding site, bioRxiv, 2020.04.17.047456 (2020).

41. J. Duan, J. Li, G. L. Chen, Y. Ge, J. Liu, K. Xie, X. Peng, W. Zhou, J. Zhong, Y. Zhang, J. Xu, C. Xue, B. Liang, L. Zhu, W. Liu, C. Zhang, X. L. Tian, J. Wang, D. E. Clapham, B. Zeng, Z. Li, J. Zhang, Cryo-EM structure of TRPC5 at 2.8-Å resolution reveals unique and conserved structural elements essential for channel function, Sci. Adv. 5, eaaw7935 (2019).

42. N. R. Gavva, J. J. S. Treanor, A. Garami, L. Fang, S. Surapaneni, A. Akrami, F. Alvarez, A. Bak, M. Darling, A. Gore, G. R. Jang, J. P. Kesslak, L. Ni, M. H. Norman, G. Palluconi, M. J. Rose, M. Salfi, E. Tan, A. A. Romanovsky, C. Banfield, G. Davar, Pharmacological blockade of the vanilloid receptor TRPV1 elicits marked hyperthermia in humans, Pain 136, 202–210 (2008).

43. A. J. Mayorga, C. M. Flores, J. J. Trudeau, J. A. Moyer, K. Shalayda, M. Dale, M. E. Frustaci, N. Katz, P. Manitpisitkul, R. Treister, S. Ratcliffe, G. Romano, A randomized study to evaluate the analgesic efficacy of a single dose of the TRPV1 antagonist mavatrep in patients with osteoarthritis, Scand. J. Pain 17, 134–143 (2017).

44. I. Sevastou, E. Kaffe, M.-A. Mouratis, V. Aidinis, Lysoglycerophospholipids in chronic inflammatory disorders: The PLA2/LPC and ATX/LPA axes, Biochim. Biophys. Acta - Mol. Cell Biol. Lipids 1831, 42–60 (2013).

45. W. Zhang, G. Sun, D. Aitken, S. Likhodii, M. Liu, G. Martin, A. Furey, E. Randell, P. Rahman, G. Jones, G. Zhai, Lysophosphatidylcholines to phosphatidylcholines ratio predicts advanced knee osteoarthritis, Rheumatol. (United Kingdom) 55, 1566–1574 (2016).

46. P. Caboni, B. Liori, A. Kumar, M. L. Santoru, S. Asthana, E. Pieroni, A. Fais, B. Era, E. Cacace, V. Ruggiero, L. Atzori, R. H. Barton, Ed. Metabolomics Analysis and Modeling Suggest a Lysophosphocholines-PAF Receptor Interaction in Fibromyalgia, PLoS One 9, e107626 (2014).

47. R. A. Rabini, R. Galassi, P. Fumelli, N. Dousset, M. L. Solera, P. Valdiguie, G. Curatola, G. Ferretti, M. Taus, L. Mazzanti, Reduced Na+-K+-ATPase activity and plasma lysophosphatidylcholine concentrations in diabetic patients, Diabetes 43, 915–919 (1994).

48. P. Liu, J. Duan, P. Wang, D. Qian, J. Guo, E. Shang, S. Su, Y. Tang, Z. Tang, Biomarkers of primary dysmenorrhea and herbal formula intervention: an exploratory metabonomics study of blood plasma and urine., Mol. Biosyst. 9, 77–87 (2013).

49. K. Hayakawa, M. Kurano, J. Ohya, T. Oichi, K. Kano, M. Nishikawa, B. Uranbileg, K. Kuwajima, M. Sumitani, S. Tanaka, J. Aoki, Y. Yatomi, H. Chikuda, Lysophosphatidic acids and their substrate lysophospholipids in cerebrospinal fluid as objective biomarkers for evaluating the severity of lumbar spinal stenosis, Sci. Rep. 9 (2019), doi: 10.1038/s41598-019-45742-7.

50. A. Riccio, Y. Li, J. Moon, K. S. Kim, K. S. Smith, U. Rudolph, S. Gapon, G. L. Yao, E. Tsvetkov, S. J. Rodig, A. Van’t Veer, E. G. Meloni, W. A. Carlezon, V. Y. Bolshakov, D. E. Clapham, Essential Role for TRPC5 in Amygdala Function and Fear-Related Behavior, Cell 137, 761–772 (2009).

51. B. J. Kolber, M. C. Montana, Y. Carrasquillo, J. Xu, S. F. Heinemann, L. J. Muglia, R. W. Gereau Iv, R. Gereau Iv, Activation of metabotropic glutamate receptor 5 in the amygdala modulates pain-like behavior 1/2 (ERK1/2), J Neurosci. June 16, 8203–8213 (2010).

52. L. W. Crock, B. J. Kolber, C. D. Morgan, K. E. Sadler, S. K. Vogt, M. R. Bruchas, R. W. Gereau, Central amygdala metabotropic glutamate receptor 5 in the modulation of visceral pain., J. Neurosci. 32, 14217–26 (2012).

53. K. D. Phelan, U. T. Shwe, J. Abramowitz, H. Wu, S. W. Rhee, M. D. Howell, P. E. Gottschall, M. Freichel, V. Flockerzi, L. Birnbaumer, F. Zheng, Canonical transient receptor channel 5 (TRPC5) and TRPC1/4 contribute to seizure and excitotoxicity by distinct cellular mechanisms, Mol. Pharmacol. 83, 429–438 (2013).

54. C. Pászty, C. M. Brion, E. Manci, H. E. Witkowska, M. E. Stevens, N. Mohandas, E. M. Rubin, Transgenic knockout mice with exclusively human sickle hemoglobin and sickle cell disease, Science (80-.). 278, 876–878 (1997).

55. A. D. Weyer, C. L. O’Hara, C. L. Stucky, Amplified Mechanically Gated Currents in Distinct Subsets of Myelinated Sensory Neurons following In Vivo Inflammation of Skin and Muscle, J. Neurosci. 35, 9456–9462 (2015).

56. A. Cowie, C. Stucky, A Mouse Model of Postoperative Pain, BIO-PROTOCOL 9 (2019), doi: 10.21769/bioprotoc.3140.

57. I. Decosterd, C. J. Woolf, Spared nerve injury: an animal model of persistent peripheral neuropathic pain, Pain 87, 149–158 (2000).

58. R. E. Sorge, L. J. Martin, K. A. Isbester, S. G. Sotocinal, S. Rosen, A. H. Tuttle, J. S. Wieskopf, E. L. Acland, A. Dokova, B. Kadoura, P. Leger, J. C. S. Mapplebeck, M. McPhail, A. Delaney, G. Wigerblad, A. P. Schumann, T. Quinn, J. Frasnelli, C. I. Svensson, W. F. Sternberg, J. S. Mogil, Olfactory exposure to males, including men, causes stress and related analgesia in rodents., Nat. Methods 11, 629–32 (2014).

59. K. E. Sadler, K. J. Zappia, C. L. O’Hara, S. N. Langer, A. D. Weyer, C. A. Hillery, C. L. Stucky, Chemokine (c-c motif) receptor 2 mediates mechanical and cold hypersensitivity in sickle cell disease mice, Pain 159, 1652–63 (2018).

60. S. R. Chaplan, F. W. Bach, J. W. Pogrel, J. M. Chung, T. L. Yaksh, Quantitative assessment of tactile allodynia in the rat paw., J. Neurosci. Methods 53, 55–63 (1994).

61. W. J. Dixon, The Up-and-Down Method for Small Samples, J. Am. Stat. Assoc. 60, 967 (1965).

62. Q. Hogan, D. Sapunar, K. Modric-Jednacak, J. B. McCallum, Detection of neuropathic pain in a rat model of peripheral nerve injury, Anesthesiology 101, 476–487 (2004).

63. A. M. Cowie, F. Moehring, C. O’Hara, C. L. Stucky, Optogenetic Inhibition of CGRPα Sensory Neurons Reveals Their Distinct Roles in Neuropathic and Incisional Pain, J. Neurosci. 38, 5807–5825 (2018).

64. T. King, L. Vera-Portocarrero, T. Gutierrez, T. W. Vanderah, G. Dussor, J. Lai, H. L. Fields, F. Porreca, Unmasking the tonic-aversive state in neuropathic pain, Nat. Neurosci. 12, 1364–1366 (2009).

65. R. B. Griggs, M. T. Bardo, B. K. Taylor, Gabapentin alleviates affective pain after traumatic nerve injury., Neuroreport 26, 522–7 (2015).

66. C. L. Cunningham, N. K. Ferree, M. A. Howard, Apparatus bias and place conditioning with ethanol in mice, Psychopharmacology (Berl). 170, 409–422 (2003).

67. P. W. Reeh, Sensory receptors in a mammalian skin – nerve in vitro preparation, Prog. Brain Res. 74, 271–276 (1988).

68. F. Moehring, A. M. Cowie, A. D. Menzel, A. D. Weyer, M. Grzybowski, T. Arzua, A. M. Geurts, O. Palygin, C. L. Stucky, Keratinocytes mediate innocuous and noxious touch via ATP-P2×4 signaling, Elife 7 (2018), doi: 10.7554/eLife.31684.

69. M. Koltzenburg, C. L. Stucky, G. R. Lewin, Receptive Properties of Mouse Sensory Neurons Innervating Hairy Skin, J. Neurophysiol. 78, 1841–1850 (1997).

70. J. Sostare, R. Di Guida, J. Kirwan, K. Chalal, E. Palmer, W. B. Dunn, M. R. Viant, Comparison of modified Matyash method to conventional solvent systems for polar metabolite and lipid extractions, Anal. Chim. Acta 1037, 301–315 (2018).

71. Y. Akbulut, H. J. Gaunt, K. Muraki, M. J. Ludlow, M. S. Amer, A. Bruns, N. S. Vasudev, L. Radtke, M. Willot, S. Hahn, T. Seitz, S. Ziegler, M. Christmann, D. J. Beech, H. Waldmann, Englerin a is a potent and selective activator of TRPC4 and TRPC5 calcium channels, Angew. Chemie - Int. Ed. 54, 3787–3791 (2015).

